# Dietary Serine Deprivation Impacts H3K27 and H3K4 Methyl Epigenomes to Impede Head and Neck Cancer Cell Plasticity and Tumor Growth

**DOI:** 10.1101/2025.10.21.683420

**Authors:** Stacy A. Jankowski, Lina Kroehling, Emily R. Fisher, Nina C. Hardy, Bach-Cuc Nguyen, Manish V. Bais, Xaralabos Varelas, Stefano Monti, Maria A. Kukuruzinska

**Author notes:** These authors contributed equally to this work (Stacy A. Jankowski, Lina Kroehling).

## Abstract

Oral squamous cell carcinoma (OSCC) is an aggressive head and neck malignancy characterized by high morbidity, therapeutic resistance and intratumoral heterogeneity driven by plastic cell states. Given that metabolic inputs can shape cell identities via epigenetic mechanisms, we investigated how metabolism of a non-essential amino acid, serine, affects histone modifications with key roles in cell plasticity: H3K27me3, which represses differentiation genes, and H3K4me3 which activates stemness and epithelial-to-mesenchymal transition (EMT) genes. Using a panel of human OSCC patient-derived cell lines and an orthotopic murine isograft model, we show that OSCC cells depend on exogenous serine for proliferation. Dietary serine deprivation induced de novo serine synthesis with a concomitant increase in α-ketoglutarate (αKG), a cofactor for KDM6B and KDM5A/B demethylases of H3K27me3 and H3K4me3, respectively. RNA-seq-derived serine deprivation gene signatures revealed activation of keratinization program and suppression of EMT and proliferation genes and tracked with good OSCC patient outcomes in TCGA. Furthermore, CUT & RUN profiling showed site-specific losses of H3K27me3 at differentiation genes and reduction of H3K4me3 at stemness, EMT and cell cycle genes. However, inhibition of αKG with 2-hydroxyglutarate was not sufficient to rescue cell proliferation. Instead, genome-wide analysis revealed widespread H3K27me3-H3K4me3 bivalency, with extensive transcriptional repression of proliferation and oncogenic programs. Functionally, serine deprivation impaired orthotopic tumor growth and improved the immune landscape in syngeneic mice. Our studies identify a metabolic serine–αKG–KDM-H3K27me3/H3K4me3 bivalency axis that globally reprograms OSCC chromatin as a potential therapeutic strategy to impede tumor plasticity and evolution to advanced disease.

**Implications:** This study identifies the tumor-suppressive role of dietary serine deprivation through epigenetic reprogramming of chromatin in OSCC.

## Introduction

Head and Neck Cancer (HNC) is a complex malignancy comprising multiple anatomical subsites in the upper aerodigestive track and ranking as the seventh most prevalent malignancy worldwide with substantial recurrence rates and mortality (1, 2). Oral squamous cell carcinoma (OSCC), the most common HPV-negative subsite remains a devastating and understudied malignancy with little improvement in patient survival over the past three decades (2).

Mounting evidence indicates that tumor metastasis, resistance to therapy and recurrence are associated with heterogeneous cell states induced by alterations in gene expression programs that drive dynamic changes in cell identities or cell plasticity (3). Such variations in transcriptional signatures and cell identities, are frequently driven by epigenetic changes in the chromatin landscape without altering the DNA sequence to generate subpopulations of aggressive cells with stem cell-like properties, known as <u>c</u>ancer <u>s</u>tem <u>c</u>ells (CSCs), cells with altered cell cycle networks, as well as cells with <u>e</u>pithelial-to-<u>m</u>esenchymal transition (EMT) and EMT hybrid phenotypes (3, 4). Like other malignancies, OSCC is characterized by heterogeneous cell states whose evolution from precancer lesions to metastatic disease involves dynamic alterations in cell identities or cell plasticity (5, 6).

In cancer, the epigenetic mechanisms utilized in the dysregulation of gene expression programs include changes in DNA methylation, alterations in the expression of no-coding RNAs, and chromatin remodeling via histone modifying enzymes, including regulators of histone acetylation and methylation. Among histone modifications, trimethylation of H3K27 has a central role in the suppression of differentiation programs, while trimethylation of H3K4 is associated with an open chromatin structure and expression of stemness, EMT and proliferation genes (7, 8). Additionally, H3K27me3 together with H3K4me3 can form bivalent chromatin domains, with the repressive H3K27me3 positioned proximal to H3K4me3 at the promoters of stemness, EMT and proliferation genes thus establishing a suppressed or “poised” state, which may be resolved in favor of either gene activation and cell plasticity or gene repression and anti-tumor responses (9). While chromatin bivalency is well documented in embryonic development, its role in OSCC has not been rigorously examined.

Previous reports highlighted the important contributions of cellular metabolism, including the nutrient microenvironment, to cancer evolution and response to therapy (10). Metabolites have been shown to play important roles in cell fate decisions through the epigenomic modulation of the chromatin landscape driven, in part, by alterations in histone tail modifications including H3K4me3 and H3K27me3 (11). Notably, some metabolites serve as cofactors to epigenetic modifiers that alter histone marks and redirect cell fate (12). Moreover, recent studies have shown that a non-essential amino acid, serine, plays an important role in cell fate decisions during epidermal development, and serine deprivation has emerged either as an inhibitor of tumor progression (13), or to have pro-tumorigenic effects (10). Given that OSCC develops from the mucosal oral epithelium with heterogeneous cell states and given that the signals that drive the dynamic and unstable cell phenotypes during OSCC progression are not well understood, we aimed to determine the impact of dietary serine deprivation on OSCC evolution.

Here, we show that human patient-derived OSCC cells rely on dietary serine to sustain transcriptional programs driving stemness, EMT, proliferation and metastatic traits, and that serine deprivation suppresses OSCC proliferation in vitro and orthotopic isograft tumor growth in mice. Our studies reveal a dual mechanism restraining OSCC progression: de novo serine synthesis that elevates α-ketoglutarate (αKG), a cofactor for KDM6B/JMJD3 histone demethylase, and removes the trimethyl group from H3K27 at differentiation gene suppressive loci inducing the epithelial differentiation program and concurrent H3K27me3 location to H3K4me3-marked promoters, thus establishing bivalent chromatin domains and suppressing stemness, cell cycle and EMT programs. Collectively our studies identify the serine–αKG–KDM-H3K27me3/H3K4me3 bivalency axis as a driver of epigenetic re-programming of OSCC chromatin to anti-tumorigenic state.

## Materials and Methods

### Cell lines and cell culture

Human OSCC patient-derived CAL27 (CAT#CRL-2095; RRID:CVCL_1107), SCC9 (CAT#CRL-1629; RRID:CVCL_1685), and SCC25 (CAT#CRL-1628; RRID:CVCL_1682) cells were obtained from ATCC, while HSC-3 (CAT#JCRB0623; RRID:CVCL_1288) cells were purchased from XenoTech. Murine 4MOSC1 cells were a gift from Dr. Silvio Gutkind (University of San Diego, CA)(14). All cells were grown under standard tissue culture conditions at 37°C and 5% CO_2_. CAL27, HSC-3, SCC9, and SCC25 cells were initially cultured in Dulbecco’s modified Eagle’s medium (DMEM) (Gibco) supplemented with 10% dialyzed fetal bovine serum (FBS) (Gibco) and 1% penicillin/streptomycin (Gibco). In-house made DMEM contained MEM with no L-glutamine (Gibco), 25X MEM Vitamin Solution (Gibco), 3500 mg/L D-glucose (Gibco), 4mM L-glutamine (Gibco), 50X MEM amino acids (Gibco), 1% penicillin/streptomycin (Gibco), 0.4mM L-serine (Sigma), and 0.4mM glycine (Sigma). Serine deprivation medium (-Ser/Gly) contained all the previously mentioned components for DMEM, excluding L-serine and glycine. Determination of the effects of D2HG on cell viability in complete media and Ser/Gly deprivation conditions was performed using PrestoBlue viability assay kit, as described below.

### OSCC orthotopic isograft mouse model

Male and female 20-week-old C57BL/6 mice were purchased from The Jackson Laboratory (CAT#000664; RRID:IMSR_JAX:000664) and housed under standard conditions at the animal facility at Boston University Chobanian & Avedisian School of Medicine. All experiments followed the protocol approved by the Institutional Animal Care and Use Committee (IACUC). Mice were injected with 1.0 x 10^6^ murine 4MOSC1 cells into the tongue and tumor growth was measured every other day via caliper. Experimental diets were purchased from TestDiet, with Mod TestDiet Baker AA Diet w/ 8% Fat and 15% sucrose (5WA1) used as control diet and Modified TestDiet 5WA1 w/ No Ser or Gly (5B62) serving as - Ser/Gly diet.

### Cell proliferation and viability assays

Cell proliferation was evaluated using the CyQUANT Cell Proliferation Assay (Invitrogen C7026). Cells were seeded at 10,000 cells per well in replicate 24 well plates in complete medium. After 24 hours media was replaced with fresh complete or serine deprivation media. Samples were collected at 0, 24, 48, and 72 hours, with media refreshed every 24 hours. At the indicated times, media was aspirated, and the plate was stored at -80°C until all plates had been collected. On the day of readout, plates were thawed at room temperature for one hour and 200µL of CyQUANT GR dye/lysis, prepared using manufacturer’s instructions, was added to each well. Samples were incubated for 5 minutes, and fluorescence was measured using a TriStar LB941 (Berthold) microplate reader at 450nm excitation and 535nm emission wavelengths. Phase contrast images were taken by LifeTechnologies EVOS XL.

Cell proliferation in response to 2-hydroxyglutaric acid (D2HG) treatment was assessed using PrestoBlue Cell Viability Reagent (Invitrogen, A13261). Cells were seeded at 125,000 cells/well in 6-well plates and cultured for 24 hours before switching to complete or Ser/Gly deprivation media. At treatment initiation, cells received either 3 or 10 mM of D2HG, with untreated controls included for each media condition. Cell viability was measured at 0, 24 and 48 hours by adding 10x PrestoBlue directly to each well and incubating for 1 hour at 37°C, 5% CO2. Media was then transferred to a black 96-well plate (Falcon, 353376) and fluorescence was measured on a TriStar LB941 microplate reader (Berthold) with 540 nm excitation and 600 nm emission. After each readout, original 6-well plates were washed with 1x DPBS and replenished with fresh media and treatment. The assay was performed in three independent experiments with triplicate samples.

### Metabolomics assay and analysis

Cells were seeded in 10cm^2^ in control media and allowed to attach for 24 hours before switching to either complete or serine deprivation medium. Cell numbers were determined at time of seeding, so cells were ∼70-80% confluent at time of metabolite extraction. Polar metabolites were extracted using ice cold 80% methanol and were lyophilized using SpeedVac (Savant). Dried metabolites were submitted to Beth Israel Deaconess Medical Center (BIDMC) Mass Spectrometry Core Facility for LC-MS metabolite profiling. LC-MS profiling and peak integration were performed according to BIDMC Mass Spectrometry Core published methods in Yuan et al. (15). After receiving peak intensity alignments to known metabolites, metabolites with an NA value in at least one sample were discarded, eliminating 65 of 300 metabolites. Metabolite abundance was log2 transformed and quantile normalized. Limma v 3.50.3 (16) was used for differential expression analysis with the formula *design <-model.matrix(∼0+group),* where group was the media condition. MetaboAnalyst (17, 18) was used to perform metabolite set enrichment analysis with the SMPDB database (19). Metabolites with a padj (FDR) < 0.05 with no fold change cutoff were used as input, with all metabolites detected as the background.

### qPCR

Cells were seeded in 10cm^2^ in control media and allowed to attach for 24 hours before switching to either complete or Ser/Gly deprivation medium. At the time of RNA extraction, cells were ∼70-80% confluent. RNA was isolated using QIAshredder (Qiagen 79656) and RNeasy Mini RNA extraction kit (Qiagen 74104), and 1 ug of RNA was used for cDNA synthesis using qScript cDNA SuperMix (QuantaBio 95048). cDNAs were mixed with gene-specific primers and *Power*SYBR Green PCR Master Mix (Applied Biosystems 4368577) and qPCR was performed using StepOne Software. Values were normalized to expression of J-actin. Primer sequences can be found in Table S2.

### Western blot

Cells were seeded in 10cm^2^ in control media and allowed to attach for 24 hours before switching to either complete or serine deprivation medium. Based on seeding density, cells were ∼70-80% confluent at time of protein collection. For whole cell lysates, cells were washed in PBS and then lysed in Triton-X-100/β-octylglucoside buffer containing protease and phosphatase inhibitor (Halt, ThermoScientific). Histone extraction was performed following the protocol outlined in the Histone Extraction Kit (Abcam ab113476). All protein extractions were quantified using a Pierce BCA Protein Assay Kit (ThermoFisher 23227). BSA standard concentrations and sample absorbance values were measured on a TriStar LB941 microplate reader at 540 nm. Interpolation was calculated using GraphPad Prism 9.

For immunoblot analyses, 20 μg of whole cell lysate or 10 μg histone extract were loaded on a 4-20% Mini-PROTEAN TGX Precast gel (BioRad). Following electrophoresis, proteins were transferred to a PVDF membrane for whole cell lysate or nitrocellulose membrane for histones. Membranes were incubated overnight at 4°C for primary antibodies and incubated with HRP-linked secondary antibodies for 1 hour at room temperature the following day. Antibodies were visualized using SuperSignal West Pico PLUS Chemiluminescent Substrate (ThermoFisher) and images were acquired using Bio-Rad ChemiDoc MP Imaging System. To detect multiple proteins, membranes were stripped using Restore PLUS Western Blot Stripping Buffer (ThermoScientific), tested for stripping efficacy, blocked again for 30 minutes at room temperature and then incubated overnight at 4°C with primary antibody. Bands were quantified using Fiji (ImageJ) and statistical analyses were performed in GraphPad Prism 9. The following primary antibodies used were: rabbit anti-PHGDH (CellSignalingTechnology, RRID:AB_2737030), rabbit anti-PSAT1 (Invitrogen, RRID:AB_11153426), rabbit anti-PSPH (Invitrogen, RRID:AB_11154135), rabbit anti-tri-methyl histone 3 lysine 27 H3K27me3 (CellSignalingTechnology, RRID:AB_2616029), rabbit anti-tri-methyl histone 3 lysine 4 H3K4me3 (CellSignalingTechnology, RRID:AB_561095), mouse anti-histone 3 H3 (CellSignalingTechnology, RRID:AB_2756816), rabbit anti-E-cadherin (CellSignalingTechnology, RRID:AB_2291471), mouse anti-N-cadherin (CellSignalingTechnology, RRID:AB_2798427), rabbit anti-Galectin-1 (CellSignalingTechnology, RRID:AB_2798065), rabbit anti-PAI-1/SerpinE1 (Abcam, RRID:AB_1642775), rabbit anti-β-actin (CellSignalingTechnology, RRID:AB_1903890), rabbit anti-GAPDH (CellSignalingTechnology, RRID:AB_10622025), mouse anti-GAPH (CellSignalingTechnology, RRID:AB_2756824).

### αKG assay

Cells were seeded in 10cm^2^ in control media and allowed to attach for 24 hours before switching to either complete or serine deprivation medium. Cell numbers were determined at time of seeding and when they reached ∼70-80% confluency at endpoint. Alpha-ketoglutarate concentration was calculated using the Alpha Ketoglutarate (αKG) Assay Kit (Abcam ab83431). In brief, samples were collected using the Alpha KG Assay Buffer. Cells were trypsinized and counted to determine the total cell number. Samples were then homogenized and deproteinated with PCA and neutralized with KOH. Afterward, αKG is transaminated to generate pyruvate, which converts a colorless probe to fluorescence. Fluorescence was measured using a TriStar LB941 microplate reader at 540 nm and 600 nm wavelengths. Sample fluorescence readouts are interpolated using an alpha-ketoglutarate standard and calculated as the amount of αKG per million cells using GraphPad Prism 9.

### Immunocytochemistry

Cells were seeded in Lab-Tek II 8-well chamber slides at 10,000 cells per well and allowed to attach for 24 hours before switching to either complete or serine deprivation medium and were grown for either 48 or 72 hours. At the predetermined time, cells were fixed with 4% paraformaldehyde, permeabilized with 0.1% Triton X-100 in 1X PBS, blocked with 1% BSA in 1X PBS, and stained with respective antibodies. Washes were performed between each step using 0.05% Tween 20 in 1X PBS (PBS-T). Primary antibodies were diluted in 0.1% BSA in 1X PBS and incubated overnight at 4°C. Slides were washed in PBS-T and incubated with fluorescent-conjugated secondary antibodies in 0.1% BSA in 1X PBS for 1 hour at room temperature in the dark and washed again. F-actin was visualized using AlexaFluor 488 Phalloidin (Invitrogen) dye. Nuclei were counterstained with DAPI (ThermoScientific) for 10 minutes at room temperature in the dark, washed, and all liquid was decanted from wells. After removing the media chamber, coverslips were applied using Prolong Diamond Antifade Mountant (Invitrogen). Slides were imaged with Zeiss LSM 710-Live Duo Confocal with 2-Photon Capability using Zen software. Fluorescence intensity was measured using CellProfiler with masking on DAPI. Statistical analysis was performed in GraphPad Prism 9. The following primary antibodies were used: rabbit anti-tri-methyl histone 3 lysine 27 H3K27me3 (CellSignalingTechnology, RRID:AB_2616029 and rabbit anti-tri-methyl histone 3 lysine 4 H3K4me3 (CellSignalingTechnology, RRID:AB_561095).

### CUT & RUN assay, sequencing, and analysis

HSC-3 cells were seeded in 10cm^2^ in control media and allowed to attach overnight before switching to either complete or serine deprivation medium. Cell numbers were determined at time of seeding, so cells were ∼70% confluent at time of collection. Cleavage Under Targets & Release Under Nuclease (CUT&RUN) was performed using the EpiCypher CUTANA™ ChIC/CUT&RUN Kit. In brief, cells were harvested and incubated with concanavalin A conjugated paramagnetic beads. The cell-bead solution was incubated with each respective antibody per sample,, H3K27me3 (H3K27me3 Antibody, SNAP-Certified™ for CUT&RUN and CUT&Tag, EpiCypher, RRID:AB_3665059) and H3K4me3 (H3K4me3 Antibody, SNAP-Certified™ for CUT&RUN, EpiCypher, RRID:AB_3075423), as well as positive controls for the targets of interest and negative control, CUTANA™ IgG Negative Control Antibody for CUT&RUN (RRID:AB_2923178). Antibodies were incubated with respective samples overnight at 4°C. The next day, samples were incubated with Protein A-Protein G Micrococcal Nuclease (pAG-MNase). Following binding, the pAG-MNase is activated by the addition of calcium chloride and incubated for 2 hours. After the Stop Master Mix is added, DNA is purified using SPRIselect Bead-Based Reagent (Beckman Coulter, Inc.) and eluted using TE Buffer. Quality control, library preparation, and sequencing were performed by the Dana-Farber Cancer Institute (DFCI) Molecular Biology Core Facilities (MBCF).

CUT&RUN libraries were prepared using IDT DNAseq library prep reagents on a Beckman Coulter Biomek i7 liquid handling platform from approximately 1ng of DNA according to manufacturer’s protocol and 14 cycles of PCR amplification per DFCI MBCF instruction. Completed sequencing libraries were quantified by Qubit fluorometer and Agilent TapeStation 2200. Library pooling and indexing was evaluated with shallow sequencing on an Illumina MiSeq. Subsequently, libraries were sequenced on an Illumina NovaSeq6000 targeting 20 million 150bp read pairs by the Molecular Biology Core Facilities at Dana-Farber Cancer Institute. Sequenced reads were aligned to the UCSC hg38 reference genome assembly using BWA (v0.7.17)(20).

After receiving sequenced reads from DFCI MCBF, macs3 v 3.0.0a6 (20) was used to call broad peaks for each sample using *-B -q 0.01 --keep-dup 1 --broad -g hs -f BAMPE --extsize 146 --nomodel* on the unique deduplicated and sorted bam files using the IgG as an input control file. Bam files were normalized to bigwig format for visualization using a scale factor determined by average band intensity from four replicates of western blots compared to H3 using deeptools v3.5.1 bamCoverage on each unique sorted and deduplicated bam file. Normalized bigwig files were used for visualizations (21). My_rose.py from Whyte et al. (22) and Lovén et al. was used to stitch peaks within 4000 kb of each other using -w 4000 for the Complete H3K27me3 samples. FRiP scores were calculated for each sample by dividing the total number of fragments overlapping peaks, determined using bedtools v 2.31.0 (23) intersect, by the total number of fragments. Samples with FRiP scores <0.1 were eliminated from the analysis due to a low signal to noise ratio. For conditions with at least 2 samples remaining in the analysis, a set of consensus peaks was identified consisting of peaks common to at least two (H3K27me3) or four (H3K4me3) replicates. Deeptools was used to plot fingerprint plots and heatmaps of consensus peaks. Genome region histograms were plotted using Gviz from normalized bigwig files and the hg38 genome.

### Differential peak analysis

The union of complete and serine deprivation consensus peaks for each histone mark were used with DiffBind to identify differentially bound peaks. Dba.count was used with summits = FALSE to disable centering and shrinking regions around peak summits. For the H3K4me3 analysis, which contains samples from two batches, dba.contrast was used with treatment and batch to account for batch effects. Peak regions were overlapped with genes using the hg38 genome and the ChIPseeker package. All peaks categorized within 1kb of a promoter region were included for enrichment analysis. Peaks were ranked by fold change, and KS tests were performed with the Kegg and hallmark pathways to identify enriched genesets in each condition.

### Bulk RNA sequencing

Cells were seeded in 10 cm^2^ plates in control media and allowed to attach for 24 hours before switching to either complete or serine deprivation medium. Cell numbers were determined at time of seeding, so cells were ∼70-80% confluent at time of RNA extraction. RNA extraction was performed using the RNeasy Mini RNA extraction kit (Qiagen). RNA samples were given to the Boston University Microarray & Sequencing Core to calculate RNA integrity and perform RNA sequencing. FastQC v 0.11.7 was used to evaluate the quality of the fastq files. STAR v 2.6.0c (24) was used to align reads to the human GRCh38 genome. RSEM v 1.3.1 was used to calculate expression (25). Expression matrices were converted into summarized experiment objects in R v 4.1.1. Genes with less than 25 counts across all samples were removed. DESeq2 v 1.32.0 (26) was used for differential expression analysis with the formula ∼condition. biomaRt v 2.48.3 was used to convert ensembl id to gene names. Signatures for each condition were defined as genes with false discovery rate (FDR) q-value <= 0.05 and log2fc >= 1. hypeR v 1.9.1 (27) was used to perform gene set enrichment analysis using hyper-geometric testing with the Hallmarks compendium (28) using all genes in the DESeq output as background. HPV-negative TCGA OSCC bulk RNAseq data (29) was scored with the bulk RNA-seq derived HSC-3 and CAL27 serine deprivation and complete media signatures using GSVA v 1.40.1 (30). Jonckheere tests were used to test for monotonic association between signature scores and stage or grade. GSEA v 4.3.3 (31) was used to test for enrichment of the keratinization signature (32) in bulk RNAseq experiments.

### H&E and IF staining and imaging of FFPE mouse tongue tissue

Formalin-fixed mouse tongue tissues were embedded in paraffin by iHisto Histopathology Support. For H&E staining, sections were deparaffinized and hydrated before being stained with eosin Y phloxine (Sigma) for cytoplasmic visualization and Harris Hematoxylin (Leica) for nuclei. H&E slides were imaged using the Zeiss Axio Scan.Z1 microscope with Zen (v3.1) Blue Software. For IF staining, formalin-fixed sections were deparaffinized in graded xylene and ethanol and rinsed in deionized water. For antigen retrieval, slides were placed in Antigen Unmasking Solution (Vector Laboratories) in a pressure cooker at high pressure for 15 minutes. Samples were brought to room temperature by rinsing with deionized water and blocked in 5% goat serum in PBS and Rodent Block M (Biocare Medical). After rinsing in PBS, samples were incubated with primary antibodies overnight at 4°C. The next day, samples were washed with PBS and incubated with secondary antibodies for 1 hour at room temperature in the dark. Sections were washed with PBS before incubation with DAPI (ThermoScientific) for 10 minutes at room temperature in the dark. Coverslips were applied with Prolong Diamond Antifade Mountant (Invitrogen). IF slides were imaged with Zeiss LSM 710-Live Duo Confocal with 2-Photon Capability using Zen (v3.5) software. Fluorescence intensity was measured using CellProfiler with masking on E-cadherin staining to identify tumor regions. Statistical analysis was performed in GraphPad Prism 9.

### Statistical analysis

Statistical analyses were performed using Prism 9 (GraphPad) software. All in vitro experiments shown included three technical replicates and they were repeated with at least three biological replicates. Representative data are shown, unless otherwise indicated. Unpaired two-tailed Student’s t-test was used to calculate statistical significance between complete and serine deprivation conditions. One-way analysis of variance (ANOVA) was used to assess statistical significance in serine rescue experiments. Figure legends provide more information on specific statistical tests. The Jonckheere’s trend test was used to test for ordered association of the complete and serine deprivation signatures with grade and stage.

## Results

### Dietary serine deprivation inhibits OSCC proliferation

Comprehensive genomic and transcriptomic analyses show that head and neck cancer cell lines recapitulate the molecular, genomic, and transcriptomic landscapes of human tumor subtypes (33). To investigate serine dependency in OSCC, we utilized a panel of cell lines representing early and advanced disease: CAL27, SCC9, and SCC25 cells from primary tongue carcinomas, and HSC-3 from cervical lymph node metastasis. Cells were cultured in complete media containing serine and glycine, or Ser/Gly deprivation media (-Ser/Gly). Though previous studies show that cancer cell proliferation depends on serine rather than glycine, and that glycine-to-serine conversion is not a major source of L-serine production (34), glycine was also removed from the media to avoid confounding effects from serine-glycine interconversion.

Over 72 to 96 hours, OSCC cells showed significantly reduced proliferation under Ser/Gly deprivation conditions across all tested lines (**Fig. 1A** and **B**; **Fig. S1A** and **B**). Compared with complete media, proliferation decreased by 70% in CAL27 and 73% in HSC-3 cells over 72 hours (**Fig. 1A** and **B**), with similar reductions of 80% in SCC25 and 42% in SCC9 cells (**Fig. S1A** and **B**). Therefore, dietary Ser/Gly starvation inhibited growth across all OSCC cell lines tested.

**Figure 1.**
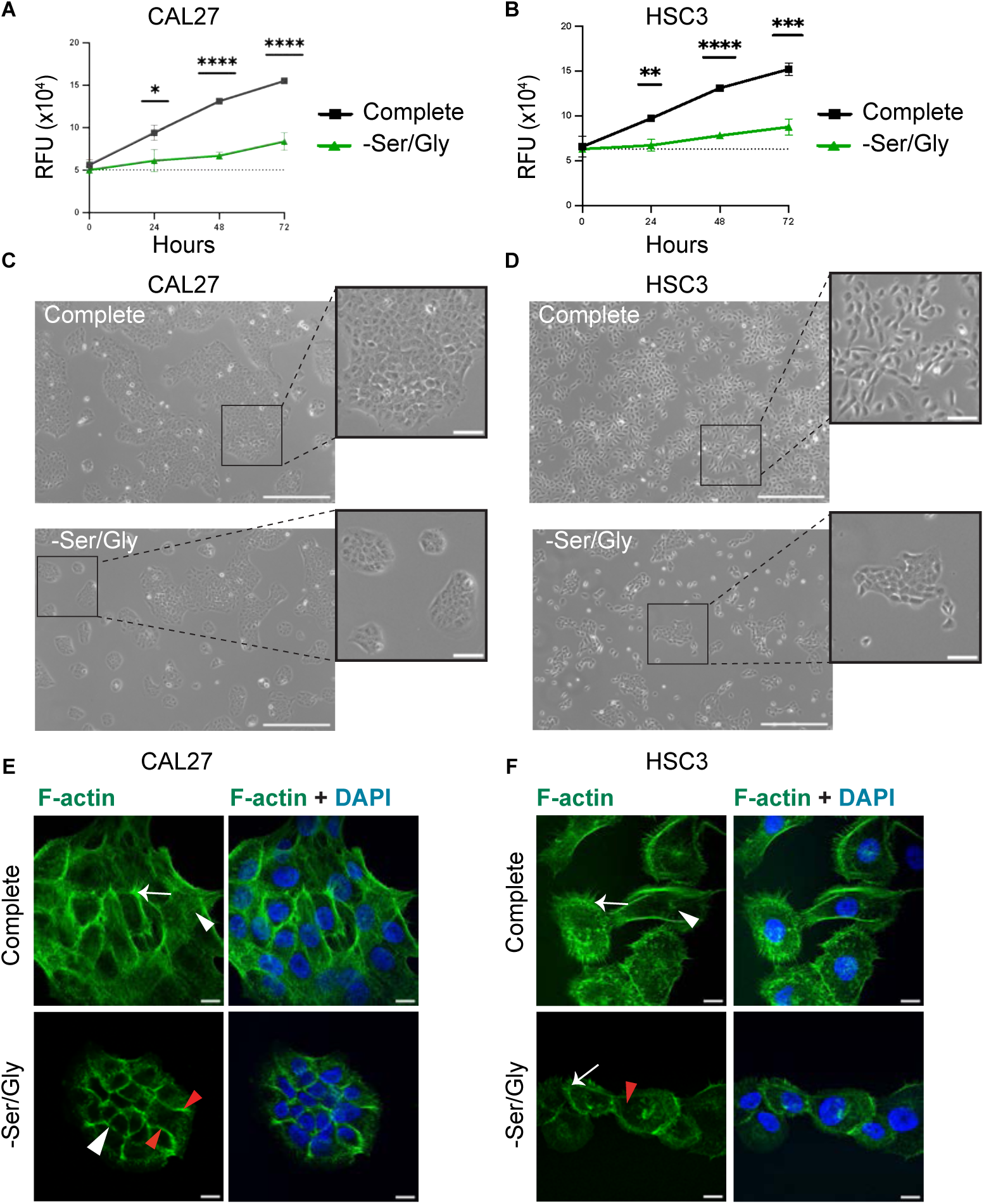
Ser/Gly deprivation inhibits proliferation of human OSCC CAL27 and HSC-3 cells and promotes an epithelial phenotype. **A,** Cell proliferation as determined by relative fluorescence units (RFU) in CAL27 cells cultured in complete medium (black) or equivalent medium lacking serine and glycine (-Ser/Gly, green). Cells showed 70% decrease in proliferation under Ser/Gly deprivation conditions. Error bars represent ±SD. Student’s unpaired t-test, n=3. **B,** Cell proliferation as determined by relative fluorescence units (RFU) in HSC-3 cells cultured in complete medium (black) or equivalent Ser/Gly deprivation medium (-Ser/Gly, green). Cells showed 73% decrease in proliferation. Error bars represent ± SD. Student’s unpaired t-test, n=3. **C,** Phase contrast imaging of CAL27 cells after 72 hours of growth in complete or -Ser/Gly media. Scale bar, 500μm; inset scale bar, 100 μm. **D,** Phase contrast imaging of HSC-3 cells after 72 hours of growth in complete or -Ser/Gly media. Scale bar, 500 μm; inset scale bar, 100 μm. **E,** Immunofluorescence imaging of F-actin in CAL27 cells grown in complete or Ser/Gly deprivation media for 72 hours. White arrowheads indicate F-actin stress fibers (top). Red arrows indicate cortical actin (bottom). Full white arrows indicate filopodia. Scale bar, 10 μm. **F,** Immunofluorescence imaging of F-actin in HSC-3 cells grown in complete or -Ser/Gly media for 72 hours. White arrowheads indicate F-actin stress fibers (top). Red arrows indicate cortical actin (bottom). Full arrows indicate filopodia. Scale bar, 10 μm.

Under Ser/Gly deprivation conditions, OSCC cells also showed altered growth patterns and changes in morphology (**Fig. 1C** and **D**; **Fig. S1C** and **D**). More aggressive oral epithelial neoplasms often display mesenchymal morphology (5). Consistent with this, HSC-3 and SCC9 cells appeared elongated in complete media but shifted toward cuboidal, epithelial-like clustering under -Ser/Gly conditions (**Fig. 1D**; **Fig. S1D**). CAL27 and SCC25 cells displayed characteristic epithelial morphology and clustered growth in complete media, which was maintained under Ser/Gly deprivation conditions (**Fig. 1C**; **Fig. S1C**).

Although CAL27 cells did not display major morphological changes under Ser/Gly restriction, immunofluorescence revealed fewer stress fibers, more organized cortical F-actin, and reduced filopodia, which contribute to cell migration, compared with complete media (**Fig. 1E**, arrowheads and arrows). Imaging HSC-3 cells showed a similar shift, with increased peripheral F-actin organization and diminished filopodia under Ser/Gly deprivation (**Fig. 1F**, arrowheads and arrows, respectively). Since mesenchymal-like cells typically display F-actin stress fibers, while epithelial cells show cortical actin organization, this redistribution suggests acquisition of differentiated epithelial traits in both cell lines following exogenous serine deprivation (35).

To confirm that Ser/Gly deprivation primarily affected the intended amino acids, we performed global metabolomics in HSC-3 cells (**Fig. S2**). Among the standard 20 amino acids, only serine and glycine were significantly altered, while changes in the three amino acids most closely related to serine and glycine, glutamate, cysteine and aspartate, were not significant (**Fig. S2A**). Of the 300+ metabolites identified under Ser/Gly deprivation, only 17 were downregulated and 15 were upregulated (**Fig. S2B** and **C**). Small Molecule Pathway Database (SMPDB) and Kyoto Encyclopedia of Genes and Genomes (KEGG) enrichment analysis confirmed that the altered metabolites included mainly intermediates in pathways converging on serine synthesis including cysteine and methionine metabolism, which rely on one-carbon units generated during serine-to-glycine conversion. Taurine and hypotaurine metabolism, associated with antioxidant protection, also increased (**Fig. S2D**) (36). Together, these data suggest that Ser/Gly deprivation primarily targeted serine and glycine metabolism, with few effects on adjacent metabolic pathways.

### OSCC cells preferentially utilize exogenous serine

Because serine and glycine can be synthesized de novo, we next investigated whether Ser/Gly deprivation induced upregulation of the endogenous serine synthesis pathway (SSP). In this pathway, glycolysis-derived 3-phosphoglycerate (3-PG) is converted to serine through sequential activity of phosphoglycerate dehydrogenase (PHGDH), phosphoserine aminotransferase (PSAT1), and phosphoserine phosphatase (PSPH), generating3-phosphohydroxypyruvate (3-PHP), 3-phospho-serine (3-P-Ser), and serine, respectively. Furthermore, serine can be converted to glycine via the bidirectional enzyme serine hydroxymethyl transferase (SHMT1/2) (**Fig. 2A**).

**Figure 2.**
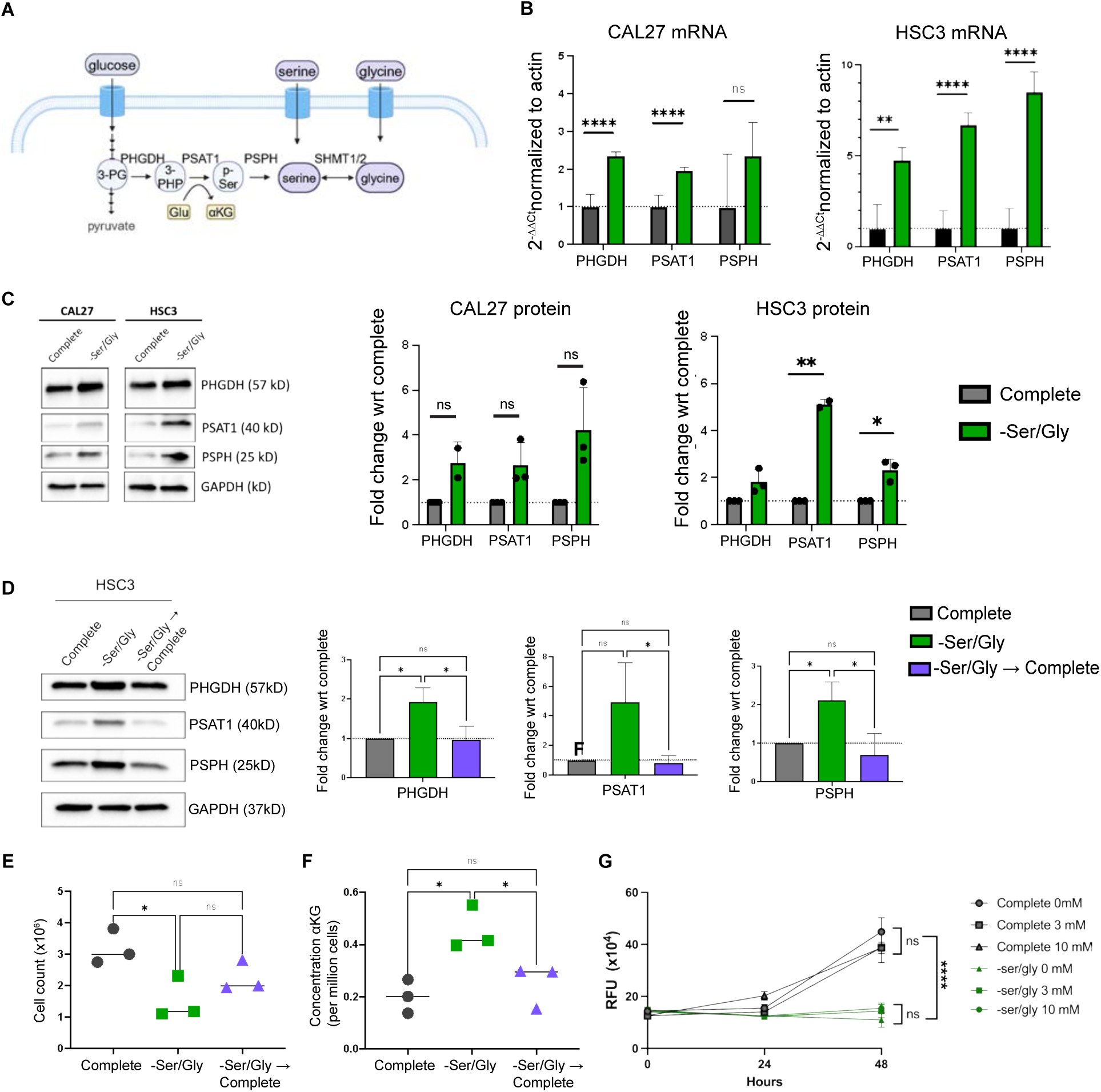
Ser/Gly deprivation drives increased expression of serine synthesis pathway enzymes and of the intermediate metabolite αKG steady-state levels. **A,** Schematic of endogenous serine synthesis pathway. Serine and glycine can be taken up from the environment through amino acid transporters or produced from the biosynthetic pathway branched from glycolysis. **B,** qRT-PCR analysis of steady-state RNA levels of serine synthesis pathway enzymes in CAL27 and HSC-3 cells grown in complete or Ser/Gly deprivation media for 48 hours. Normalization to actin using the 2^-ΔΔ^Ct method. Error bars represent ±SD. Student’s unpaired t-test, n=4. **C,** Representative immunoblot and quantification of three independent biological replicates of SSP enzymes in CAL27 and HSC-3 cells grown in complete or Ser/Gly deprivation media for 48 hours. Full blots are included as raw data files. Error bars represent ±SD. Student’s unpaired t-test, n=3. **D,** Representative immunoblot and quantification of three independent biological replicates of steady-state protein levels of SSP enzymes in HSC-3 cells grown in complete media for 72 hours (complete), Ser/Gly deprivation media for 72 hours (-Ser/Gly), or for 48 hours in Ser/Gly deprivation media followed by 24 hours in complete media (-Ser/Gly → complete). Full blots are included as raw data files. Error bars represent ± SD. Student’s unpaired t-test, n=3. **E,** Cell count per million cells of HSC-3 cells grown in complete media for 72 hours (complete), Ser/Gly deprivation media for 72 hours (-Ser/Gly), or for 48 hours in Ser/Gly deprivation media followed by 24 hours in complete media, (-Ser/Gly → complete). Error bars represent ± SD. Student’s unpaired t-test, n=3. **F,** Concentration of αKG per million cells in HSC-3 cells grown in complete media for 72 hours (complete), Ser/Gly deprivation media for 72 hours (-Ser/Gly), or for 48 hours in Ser/Gly deprivation media followed by 24 hours in complete media (-Ser/Gly → complete). Error bars represent ± SD. Student’s unpaired t-test, n=3. **G,** Quantification of changes in proliferation of HSC-3 cells grown in complete and Ser/Gly deprivation media in response to inhibition of αKG with 3 mM and 10 mM D-2HG as measured by PrestoBlue fluorescence assay. Error bars represent ± SD, student unpaired t-test, n=3 per growth condition, per D-2HG concentration, and per time point for each of the three biological replicates.

In response to Ser/Gly deprivation OSCC cells increased steady-state mRNA (**Fig. 2B**) and protein levels (**Fig. 2C**; **Fig. S2**) of PHGDH, PSAT1, and PSPH although to varying extents. PHGDH, which catalyzes the first committed and rate-limiting SSP step, has been shown to function as either anti-or pro-tumorigenic, and its modest increase in steady-state levels following serine deprivation may reflect tighter regulation of its abundance (37). In contrast, PSAT1, the second rate-limiting enzyme, was prominently increased, especially in HSC-3 cells. Consistent with prior reports that extracellular serine withdrawal increases SSP enzyme expression and pathway activity at transcriptional and translational levels (37, 38), returning Ser/Gly-deprived cells to complete media reduced steady-state levels of all three SSP enzymes (**Fig. 2D**).Together, these data suggest that OSCC cells upregulate the SSP to compensate for loss of exogenous serine.

PSAT1 is responsible for converting a glutamate to αKG (**Fig. 2A**), a tricarboxylic acid (TCA) cycle intermediate that regulates energy metabolism, and chromatin states, with downstream effects that limit proliferation and impact both lineage-specific gene expression and cell fate programs (12). Because metastatic HSC-3 cells displayed the most robust increase in PSAT1 under Ser/Gly deprivation (**Fig. 2C** and **D**), coinciding with reduced cell number (**Fig. 2E**), we quantified αKG levels. Indeed, αKG concentration per million cells increased concomitantly with PSAT1 under Ser/Gly deprivation conditions (**Fig. 2F**). Conversely, when exogenous Ser/Gly was added back to the media, aKG levels decreased, correlating with reduced PSAT1 steady-state levels and increased cell numbers (**Fig. 2E** and **2F**). Therefore, αKG levels tracked with changes in total number of cells in Ser/Gly deprivation conditions, with increased levels in Ser/Gly deprivation and decreased abundance once total cell number began to recover after switch to complete media (**Fig. 2E** and **2F**). These results demonstrate that OSCC cells preferentially utilize exogenous serine when available but activate the endogenous SSP under Ser/Gly deprivation with concomitant αKG production.

The JumonjiC-domain (JmjC) family is composed of lysine demethylases (KDM) that modify histone 3 (39), including JMJD3/KDM6B, which demethylates H3K27me3, a repressive chromatin mark that silences epithelial differentiation genes (8, 40). To test whether the observed increases in αKG levels under Ser/Gly deprivation impacted OSCC cell proliferation, we treated HSC-3 cells in complete or Ser/Gly deprivation media with 2-hydroxyglutarate enantiomer D (D2HG), a competitive αKG antagonist that inhibits its role as a co-factor for KDM6B and KDM5A/B, demethylases of H3K27me3 and H3K4me3, respectively. In complete media, with little αKG generated from the SSP pathway, 3 mM or 10 mM D2HG had no effect on proliferation, indicating that these concentrations were not toxic, and that TCA-derived αKG did not mediate a detectible proliferative effect. Likewise, D2HG did not alter proliferation under Ser/Gly deprivation, indicating that the increased αKG abundance from the SSP was not responsible for inhibition of cell proliferation (**Fig. 2G**). These results suggest that an additional αKG-independent anti-proliferative mechanism operates under dietary Ser/Gly deprivation.

### Enrichment analysis of the serine deprivation signature reveals diminished cell plasticity and increased cellular differentiation

We next investigated the effects of Ser/Gly deprivation on global transcript levels in CAL27 and HSC-3 cells using bulk RNA sequencing (RNA-seq). Both cell lines showed significant differential gene expression under Ser/Gly deprivation compared with complete media (**Fig. 3A** and **B**; **Fig. S3A** and **B**). Consistent with the greater morphological shift observed in HSC-3 cells (**Fig. 1C** and **D**), HSC-3 cells showed more transcriptional changes, with 581 upregulated and 346 downregulated genes, compared with 176 upregulated and 88 downregulated genes in CAL27 cells, using a log2FC cutoff of ±1 and FDR < 0.05 (**Fig. 3A** and **B**). *PHGDH, PSAT1,* and *PSPH* were significantly upregulated in both cell lines, further validating SSP induction under Ser/Gly deprivation (**Fig. 3A** and **B**; **Table S1**).

**Figure 3.**
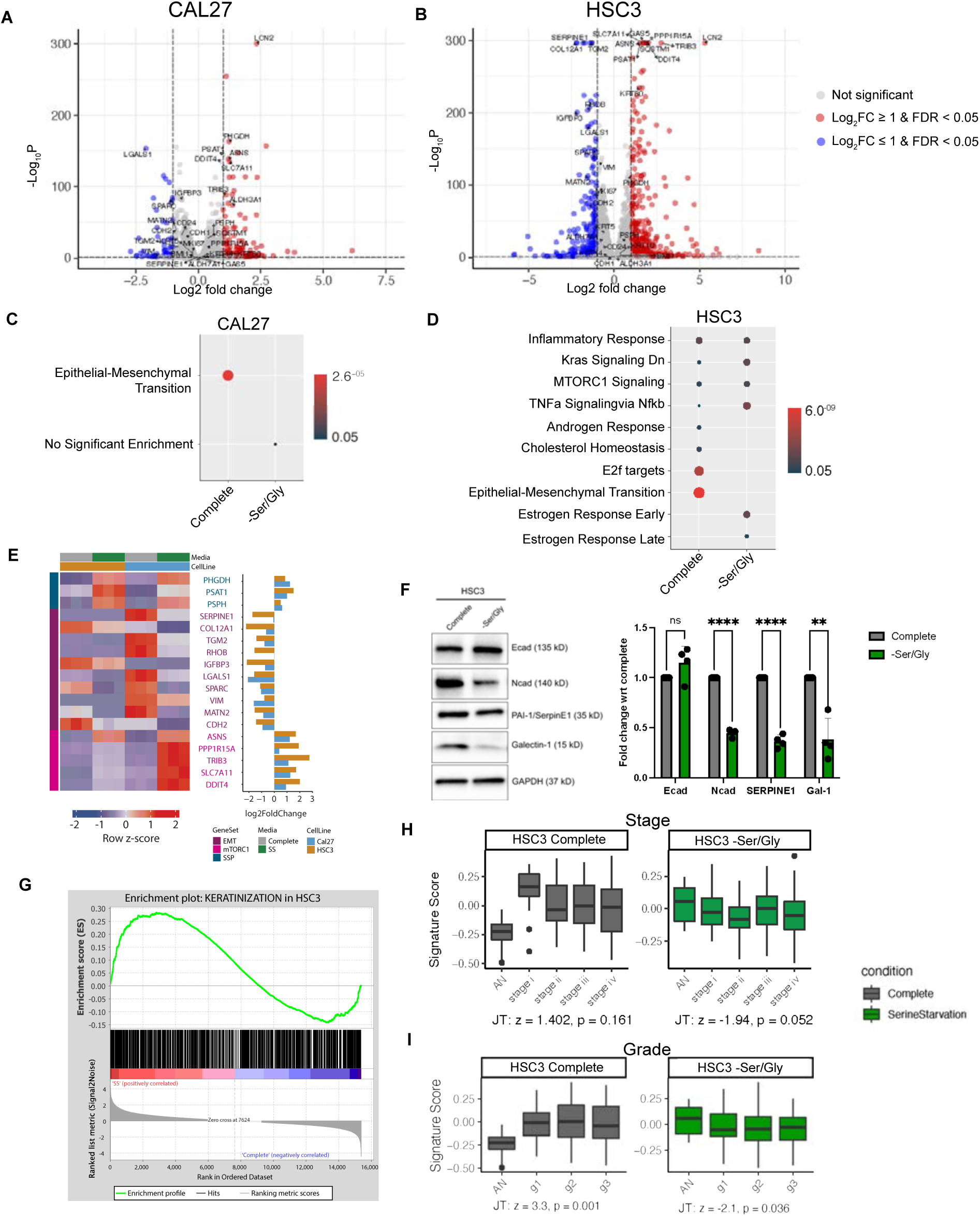
Gene set enrichment under Ser/Gly deprivation reveals diminished cell plasticity. **A** and **B,** Volcano plots highlighting top 50 most highly changed genes with FDR < 0.05 for both Log_2_-fold changes ≤ 1 (**blue**) and ≥ 1 (**red**) in bulk RNA-seq expression in CAL27 and HSC-3 cells grown in complete or Ser/Gly deprivation media for 48 hours. **C** and **D,** Gene Set Enrichment Analysis of Complete and Ser/Gly deprivation signatures derived from bulk RNA-seq with Molecular Signature Database (MsigDB) hallmark gene sets for CAL27 (**C**) and HSC-3 (**D**). Dot sizes indicate numbers of overlapping genes between signatures and gene sets, and colors indicate FDR. **E,** Heatmap of bulk RNA-seq genexpression, along with the Log_2_-fold change of select genes for CAL27 and HSC-3 cells, with three independent biological replicates per condition. **F,** Immunoblot validation and quantification of three independent biological replicates of select target genes confirming changes in gene expression in HSC-3 cells grown in Ser/Gly deprivation media for 48 hours. Student’s unpaired t-test, n=4. **G,** GSEA enrichment plot of HSC-3 RNA-seq gene signatures with squamous epithelial cell-keratinization program from the Human Protein Atlas gene set, with Normalized Enrichment Score (NES) of 1.5. **H** and **I,** HSC-3-derived RNA-seq gene signatures were projected on The Cancer Genome Atlas (TCGA) OSCC RNA-seq database with known stage and grade and used to analyze the HSC-3 Ser/Gly deprivation gene signature association with tumor progression. The Jonckheere–Terpstra (JT) test was used to determine monotonically increasing or decreasing relationships, with JT positive value in complete media indicating that signature was increasing with stage and/or grade, whereas a negative JT value for growth in Ser/Gly deprivation media indicating that signature was decreasing with increased stage/grade.

Molecular Signature Database (MSigDB) hallmark enrichment (28) identified EMT as the most significantly altered gene expression program between complete and Ser/Gly deprivation conditions (**Fig. 3C** and **D**). This was a striking finding given that EMT and partial EMT phenotypes mark aggressive OSCC subpopulations (4, 5) frequently associated with metastases and therapeutic resistance (2,3). For CAL27 cells, EMT was the only significant hallmark enriched in complete media that was strongly reduced under-Ser/Gly deprivation conditions (**Fig. 3C**). In HSC-3 cells, EMT was also the most significantly altered hallmark (**Fig. 3D**). Target hallmark *E2F* gene expression was also greatly reduced in HSC-3 cells, with downregulation of commonly overexpressed cell cycle genes *CDC20*, *CDKN3*, *CDCA3*, *CDCA8*, and *CDK1*, as well as decreased expression in proliferation marker protein *MKI67*. *mTORC1* signaling and *TNFa* signaling via *NFkb* were also altered in HSC-3 cells, with the latter linked to epigenetic regulation of histone modifications (12). Further analysis showed upregulation of genes associated with growth suppression and radio sensitization, including *PPP1R15A*, which encodes growth arrest and DNA-damage-inducible protein 34 (GADD34) (41) and *ASNS*, which directs metabolites away from activating *mTORC1*. Notably, *LNC2*, which encodes glycoprotein lipocalin-2, a metabolic stress sensor with functions in energy metabolism and inflammation was the most significantly upregulated gene in both CAL27 and HSC-3 cell lines (41).

Analysis of changes in EMT-specific genes revealed significant downregulation of markers associated with poor OSCC clinical outcomes (**Fig. 3E**; **Table S3**). These included *SerpinE1*, *TGM2*, *LGALS1* (42), and *CDH2* (N-cadherin), which are mesenchymal markers upregulated during EMT. Immunoblot validation confirmed reduced steady-state levels of these EMT markers under Ser/Gly deprivation conditions (**Fig. 3F**) along with a reciprocal change in E- and N-cadherins. Together, these results suggest that Ser/Gly deprivation transcriptionally suppresses aggressive OSCC phenotypes.

In addition to reduced EMT gene expressions, we assessed keratinocyte differentiation using Gene Set Enrichment Analysis (GSEA) (31). The Human Protein Atlas Squamous Epithelial Cell-Keratinization Signature (32) was significantly enriched in both CAL27 and HSC-3 cells under Ser/Gly deprivation genes, indicating activation of keratinocyte differentiation programs (**Fig. 3G**; **Fig. S3C**). These findings suggest that Ser/Gly deprivation promotes a more differentiated OSCC phenotype consistent with the observed morphological shift from mesenchymal to more epithelial phenotypes in Ser/Gly deprivation (**Fig. 1**, **Fig. S1**).

To align RNA-seq gene signatures to human OSCC patient outcomes, we projected 311 OSCC RNA-seq samples from The Cancer Genome Atlas (TCGA) onto our gene signatures from HSC-3 and CAL27 cells grown in complete or Ser/Gly deprivation media (39). The HSC-3 complete-media signature showed a nonsignificant association with tumor stage relative to normal tissue (**Fig. 3H**, *Complete*) but was significantly associated with tumor grade (p=0.001) and aligned with advanced disease and poor outcomes (**Fig. 3I**, *Complete*). In contrast, the HSC3 Ser/Gly deprivation signature significantly aligned with normal tissue and showed no association with progressive stage (p=0.052) or grade (p=0.036), consistent with favorable patient outcomes (**Fig. 3H** and **3I**; *-Ser/Gly*). Similarly, gene signatures from more indolent CAL27 cells grown in complete media showed a nonsignificant trend with stage and significant association with increasing tumor grade (p=0.017), whereas the CAL27 Ser/Gly-depravation signature was not associated with either stage or grade (**Fig. S3D** and **E**). These data indicated that Ser/Gly deprivation gene signatures are not associated with progression to advanced OSCC in TCGA.

### Serine deprivation leads to a loss of repressive H3K27me3 from sites associated with differentiation programs

The effect of Ser/Gly deprivation on H3K27me3 steady-state levels was assessed by immunoblot analyses of total cell lysates derived from cells grown in complete and Ser/Gly deprivation media. There was a visible, but variable, decrease in the steady state levels of H3K27me3 in HSC-3 cells under Ser/Gly deprivation, with the most noted deviation apparent in replicate # 1 (**Fig. 4A**). This variation could be due to the complexity of epigenetic regulation known to involve stochastic changes and entropy states impacting multiple gene expression networks all likely influenced by numerous factors, which can result in variations among samples. This result was supported by a significant reduction in H3K27me3 immunofluorescence concomitant with increased epithelial morphology (**Fig. 4B**, *double arrows*), which accompanied the change in F-actin from stress fibers to a more organized cortical actin (**Fig. 4B**, *arrowheads*). These results suggested that Ser/Gly deprivation caused a detectable decrease in H3K27me3.

**Figure 4.**
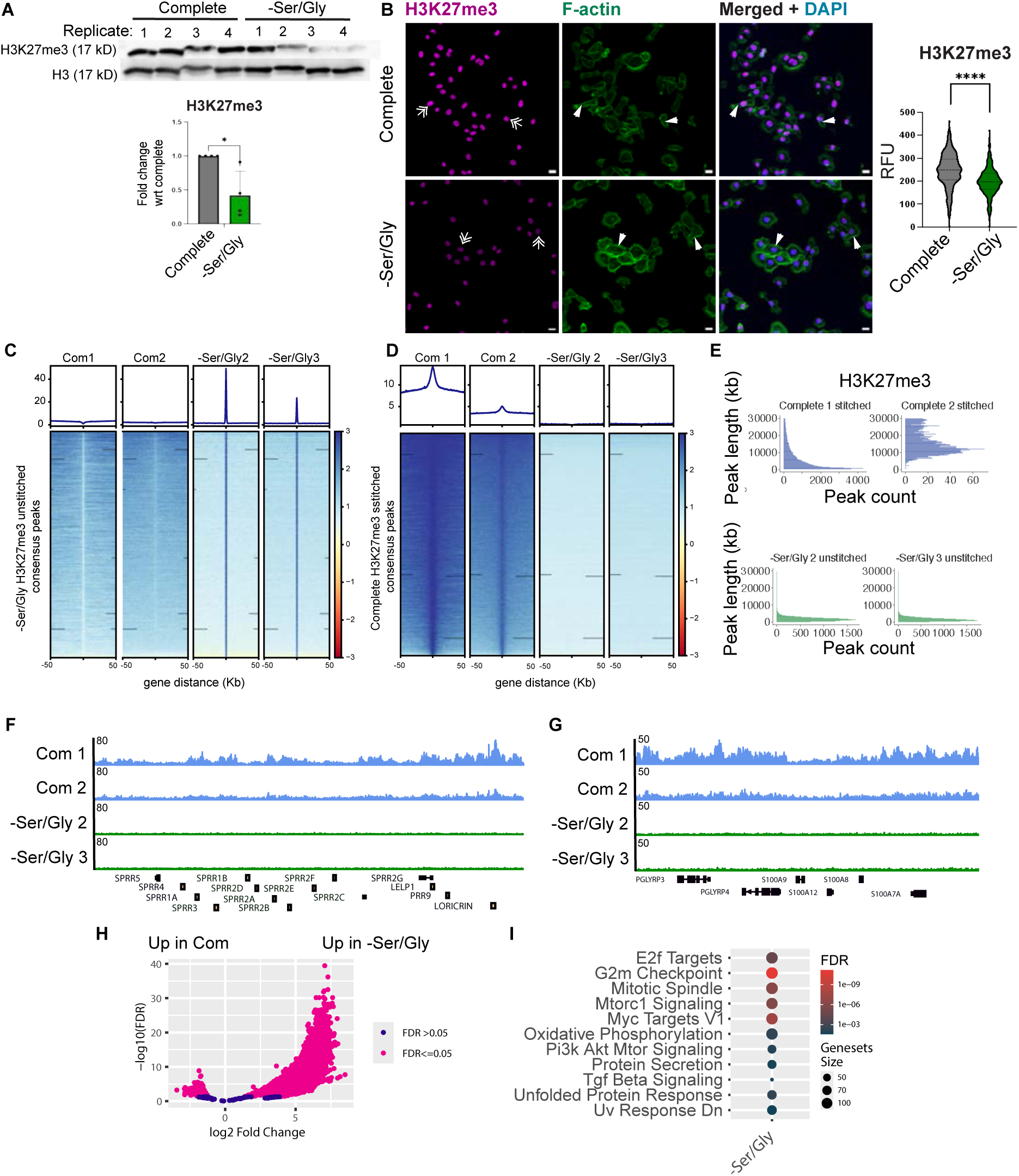
Serine deprivation leads to the reduction of repressive H3K27me3 from the differentiation genes’ loci. **A,** Representative immunoblot and quantification of histone mark H3K27me3 in four independent biological replicates of HSC-3 cells grown in complete and Ser/Gly deprivation media for 48 hours. Uncropped immunoblots are available as raw data sets. Error bars represent ± SD. Student’s unpaired t-test, n=4. **B,** Representative immunofluorescence imaging of H3K27me3 (double arrows) counterstained with DAPI and F-actin (arrowheads) in HSC-3 cells grown in complete and Ser/Gly deprivation media for 48 hours. Scale bar, 20 μm. Violin plots display a range of immunofluorescence intensities across replicates with quantification representing an average of three independent biological replicates. Student’s unpaired t-test, n=3. **C,** CUT & RUN heatmap of Ser/Gly deprivation H3K27me3 consensus peaks in HSC-3 cells grown in complete (Com 1 and 2) or Ser/Gly deprivation (-Ser/Gly 2 and 3) media for 48 hours. Region - 50Kb to +50kb around the center of the peak region is shown. **D,** CUT & RUN heatmap of Complete H3K27me3 consensus peaks in HSC-3 cells grown in complete (Com 1 and 2) or Ser/Gly deprivation (- Ser/Gly 2 and 3) media for 48 hours. Region -50Kb to +50kb around the center of the peak region is shown. **E,** Representation of stitched and unstitched peaks for H3K27me3 from HSC-3 cells grown in complete and Ser/Gly deprivation media for 48 hours. **F** and **G,** Integrative Genome Viewer (IGV) visualization of peaks spanning differentiation genes showing increased H3K27me3 coverage in complete media and reduced coverage in -Ser/Gly media at 48 hours of growth. **H,** Volcano plot showing differential peaks from H3K27me3 CUT & RUN between Ser/Gly deprivation (-Ser/Gly) and Complete (Com) media conditions. Peaks upregulated in -Ser/Gly condition are to the right of 0, and those upregulated in Complete media are to the left. Peaks significantly different between conditions (FDR <= 0.05) are in pink. **I,** KS test of mSigdbR Hallmarks pathways. Dot color corresponds to FDR, and dot size corresponds to hallmark geneset size.

To determine how Ser/Gly deprivation impacted the chromatin landscape, we investigated the changes in histone modifications in HSC-3 cells using Cleavage Under Targets & Release Under Nuclease (CUT & RUN). For these studies, cells were grown for 48 hours in either complete or Ser/Gly deprivation media and DNA was collected after targeting H3K27me3 with a specific antibody. After sequencing, replicates that had a Fraction of Reads in Peaks (FRiP) score of less than 0.1 were removed from analysis (**Table S4**). We observed that H3K27me3 presented consistently as wide peaks throughout the genome under growth in complete media conditions (**Fig. 4C** and **4E**). Given that H3K27me3 is a repressive chromatin mark and that H3K27me3-rich regions can function as super-repressors, clusters of H3K27me3-rich regions in the chromatin landscape can act either locally or via long-range chromatin interactions to silence gene expression (8). Thus, in our analyses we utilized the method of “stitching” constituent peaks as described in Cai, et al (8). Results revealed much greater peak lengths in complete media than in Ser/Gly deprivation (**Fig. 4D** and **4E**), with peaks displaying a high consensus between the two replicates (**Fig. 4D**). The consensus peaks identified in both replicates of cells grown in complete media were not detected in Ser/Gly deprivation replicates (**Fig. 4D**). The reverse was also true – peaks observed in Ser/Gly deprivation replicates were not found in the complete media replicates (**Fig. 4C**). By inspecting chromosomal peak locations using the Integrative Genome Viewer (28), we observed a decrease in H3K27me3 marks at differentiation genes including the SPRR gene family, proteins induced during differentiation (43), calprotectin, (44), and loricrin, a terminal differentiation keratinocyte protein (**Fig. 4F**). Additionally, there was a substantial decrease in H3K27me3 peaks at genes coding for the heterodimeric complex of S100A8/9, which is also a tumor suppressor in HNC (**Fig. 4G**) (44) as well as PGLYRP3, with long recognized involvement in innate immunity. These data further supported the conclusion that increased SSP activity under Ser/Gly deprivation led to the loss of H3K27me3 from its repressive sites at differentiation genes.

The observed decrease in the abundance of H3K27me3 as measured by immunoblots and immunofluorescence coincided with the loss of this histone mark from the repressive loci of selected differentiation genes, suggesting decreased levels of this repressive histone mark under Ser/Gly deprivation. Surprisingly, however, a volcano plot of genome-wide H3K27me3 peaks showed a significant (FDR≤0.05) and substantial increase in the number of upregulated H3K27me3 peaks under Ser/Gly deprivation conditions, namely 10,179 in Ser/Gly deprivation compared to 1,463 in complete media (**Fig. 4H**). Significantly, analyses of hallmark pathways associated with this increase in the H3K27me3 repressive peaks corresponded to the suppression of gene expression programs associated with E2F targets, G2-M phase transition, Myc targets, PI3K/Akt/Mtor and TGFβ signaling, shown to play key roles in OSCC tumorigenesis (2, 45)(**Fig. 4I**). This lack of alignment between immunoblots and immunofluorescence and global CUT & RUN analyses suggested that changes in the chromatin topology may underlie this discrepancy.

### Serine deprivation correlates with a decrease of H3K4me3 levels at stemness and EMT-associated genes

Given the observed loss of H3K27me3 from the selected differentiation genes loci, we sought to investigate the effects of Ser/Gly deprivation on stemness and EMT markers regulated by H3K4me3. Like H3K27me3 under Ser/Gly deprivation conditions, we observed some variation among independent replicate samples and an overall decrease in the steady-state protein levels of H3K4me3 in metastatic HSC-3 cells with replicate 1 representing the most prominent outlier for H3K4me3 (**Fig. 5A**).

**Figure 5.**
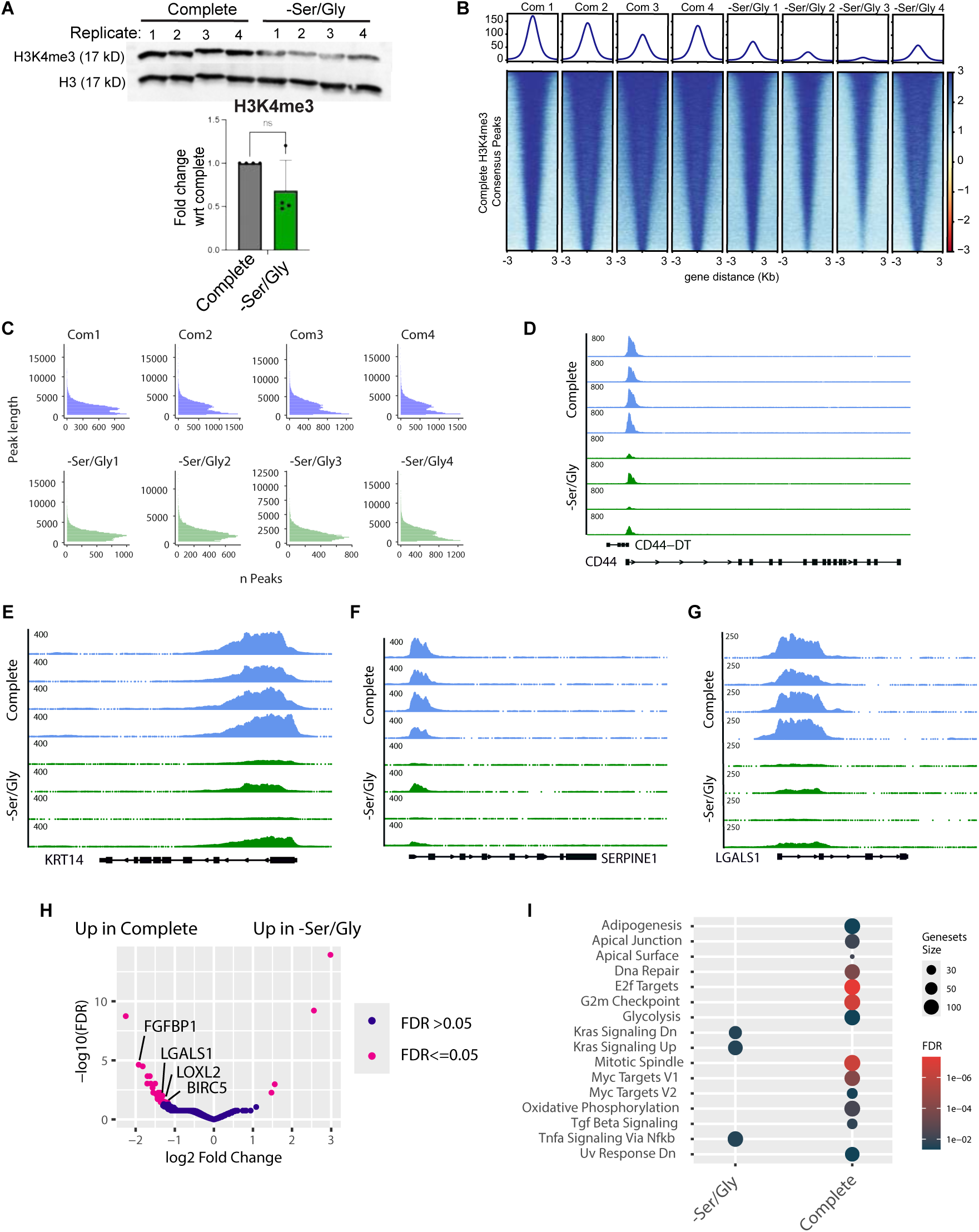
Growth of HSC-3 cells in Ser/Gly deprivation media is associated with the reduction of H3K4me3 marks. **A,** Representative immunoblot and quantification of H3K4me3 mark in four independent biological replicates of HSC-3 cells grown in complete and Ser/Gly deprivation media for 48 hours showing a 48% average decrease in Ser/Gly deprivation media compared to complete media. Uncropped immunoblots are available in raw data sets. Error bars represent ± SD. Student’s unpaired t-test, n=4. **B,** CUT & RUN sequencing heatmap of H3K4m3 peaks in HSC-3 cells grown in complete (Com 1) or Ser/Gly deprivation (-Ser/Gly 1, 2 and 3) media for 48 hours with visualization of consensus peaks significantly called in complete media sample. Region -3Kb to +3kb around the center of the peak is shown. **C,** Representation of peaks for H3K4me3 from HSC-3 cells grown in complete and Ser/Gly deprivation media for 48 hours reveals similar focused peaks in either condition. **D,** CUT & RUN visualization of peaks overlapping stemness marker CD44. Complete H3K4me3 samples are in blue, Ser/Gly deprivation (-Ser/Gly) H3K4me3 samples are in green. **E,** CUT & RUN visualization of peaks overlapping stemness marker KRT14. Complete H3K4me3 samples are in blue, Ser/Gly deprivation (-Ser/Gly) H3K4me3 samples are in green. **F,** CUT & RUN visualization of peaks overlapping EMT gene SERPINE1. Complete H3K4me3 samples are in blue, Ser/Gly deprivation (-Ser/Gly) H3K4me3 samples are in green. **G,** CUT & RUN visualization of peaks overlapping EMT gene LGALS1. Complete H3K4me3 samples are in blue, Ser/Gly deprivation (-Ser/Gly) H3K4me3 samples are in green. **H,** Volcano plot showing differential peaks from H3K4me3 CUT & RUN between Ser/Gly deprivation and complete media growth conditions at 48 hours. Peaks upregulated in Ser/Gly deprivation condition are to the right of 0, and those upregulated in Complete media are to the left. Peaks significantly different (FDR <= 0.05) between conditions are in pink. **I,** KS test of peaks ranked by FDR with pathways from MSigDB Hallmarks collection. Dot color corresponds to FDR, and dot size corresponds to the hallmark geneset size.

Next, we investigated changes in H3K4me3 in response to Ser/Gly deprivation using CUT & RUN-sequencing. After removing replicates that had a FRiP score of less than 0.1, we observed consensus peaks between cells grown in complete and Ser/Gly deprivation media (**Fig. 5B**, **Table S4**). In contrast to H3K27me3, we did not observe any wide peaks for H3K4me3 (**Fig. 5C**). Additionally, while H3K27me3 had consensus peaks specific to each condition, we found sharp peaks of H3K4me3 at the same sites but with varying amplitudes between complete and Ser/Gly deprivation conditions (**Fig. 5B** and **5C**). Specifically, under Ser/Gly deprivation conditions, we found a decrease in peaks at transcription sites of OSCC stemness markers, such as CD44 and KRT14 (**Fig. 5D** and **E**), as well as at EMT markers, such as SEPRINE1 and LGALS1 (**Fig. 5F** and **G**). These results suggested that Ser/Gly deprivation led to the reduction of H3K4me3 chromatin occupancy at the promoters of stemness and EMT genes. Thus, the loss of H3K27me3 from regions of epithelial differentiation genes (**Fig. 4**), coincided with the reduction of H3K4me3 occupancy at the promoters of stemness and EMT genes.

Global analysis of changes in H3H4me3 peaks revealed 34 peaks significantly increased in complete media, while in Ser/Gly deprivation, only 4 peaks were significantly increased (FDR<=0.05) (**Fig. 5H**). The increased peaks in the complete media were aligned with promoters of genes associated with pro-tumorigenic activities in OSCC, including *LGALS1*, *BIRC5*, *LOXL2* and *FGFBP1*, while in serine deprivation media the identified genes included *MKKS*, involved in cytokinesis and centriole duplication and a maker of HNC, miR-3189, a known tumor suppressor in many cancers, and *RASSF9*, a modulator of RAS signaling pathway with both pro- and anti-oncogenic functions, but whose role in OSCC has not been yet elucidated. The fourth peak was at the promoter of LOC107986606, although to date, its identity has not been defined. Pathway enrichment of the H3K4me3 peaks with the hallmark pathways in complete media included E2F targets, DNA repair, several cell cycle G2-M checkpoint controls, and Myc targets, among others, while in Ser/Gly deprivation conditions the most significant pathways included Kras signaling (up/down) downstream from EGFR activity, and proinflammatory TNFa signaling via NFkb, also detected in unbiased RNA-seq results (**Fig. 3D**), both absent from cells grown in complete media (**Fig. 5I**).

### Global analysis of CUT & RUN datasets reveals genome-wide H3K27me3/H3K4me3 bivalency

Together, H3K27me3 and H3K4me3 can form a bivalent chromatin structure in embryonic development, cancer and in normal differentiated tissues, which in embryonic stem cells can direct a poised state capable of resolving into either activation, silencing or lasting repression of developmental genes. In cancer, depending on the tumor type, bivalent chromatin has been shown to be aligned with cell plasticity and cancer progression (9) although epigenetic changes in chromatin topology have also been shown to restrain tumor progression (46). Given that the roles of bivalent chromatin have not been rigorously examined in OSCC, we first assessed the presence of repressive H3K27me3 marks (47) at the promoters of selected H3K4me3-associated genes. Significantly, we found H3K27me3 marks at H3K4me3 at EMT genes LGALS1 and SERPINE1 loci (**Fig. S4A** and **B**), indicating the presence of bivalent chromatin structures. This suggested that under conditions of Ser/Gly deprivation, the repressive H3K27me3 was recruited to silence H3K4me3 marked genes.

To test whether bivalent chromatin formation occured more broadly, we performed genome-wide overlap analysis of CUT & RUN sequencing data. In complete media, H3K4me3 peaks showed no overlap with H3K27me3, and vice versa (**Fig. 6A**), indicating absence of bivalency. In contrast, under Ser/Gly deprivation, H3K4me3 consensus peaks coincided with H3K27me3 peaks across the genome (**Fig. 6B**), indicating widespread bivalent domains formation. A genome-wide map revealed extensive bivalency across all human chromosomes, with increased domain density in select regions (**Fig. S4C**). These results demonstrate that Ser/Gly deprivation induces widespread H3K27me3-H3K4me3 bivalent chromatin formation that suppresses aggressive cell-state genes coincident with de-repression of differentiation programs.

**Figure 6.**
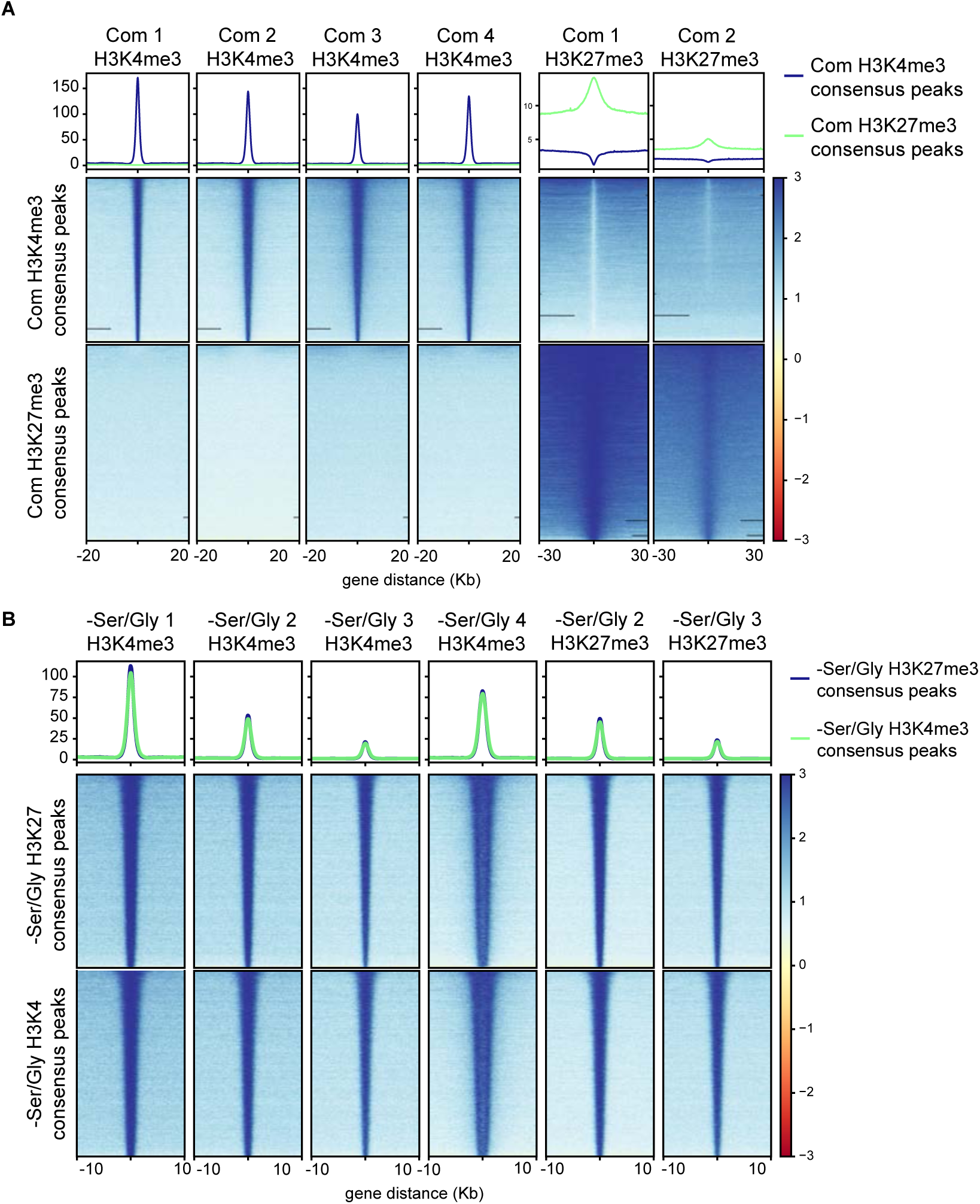
Ser/Gly deprivation induces genome-wide H3K27me3-H3K4me3 chromatin bivalency program. **A,** CUT & RUN sequencing heatmap of complete H3K4me3 consensus peaks (top row) in HSC-3 cells grown in complete media for 48 hours with H3K4me3 CUT&RUN (first four columns) or H3K27me3 CUT&RUN (last 2 columns); and Complete H3K27me3 consensus peaks (bottom row), in the same six samples. Consensus peaks shown identified in each histone mark are unique to that histone mark. Region -20Kb to +20kb around the center of the peak region is shown for the H3K4me3 samples, and -30Kb to +30kb around the center of the peak for H3K27me3 samples. **B,** CUT & RUN sequencing heatmap of Ser/Gly deprivation H3K27me3 consensus peaks (top row) in HSC-3 cells grown in Ser/Gly deprivation media for 48 hours with H3K4me3 CUT & RUN (first four columns) or H3K27me3 CUT&RUN (last 2 columns); and -Ser/Gly H3K4me3 consensus peaks (bottom row), in the same six samples. Region -10Kb to +10kb around the center of the peak region is shown. Consensus peaks shown identified in each histone mark are shared across both histone marks, indicating chromatin bivalency.

### Serine deprivation diet inhibits orthotopic OSCC tumor growth in syngeneic mice

To validate the impact of Ser/Gly deprivation on OSCC evolution in vivo, we examined orthotopic tumor growth in syngeneic C57BL/6 mice. We used the murine 4MOSC1 cell line derived from a tobacco-mimetic carcinogen, 4-nitroquinoline-1-oxide- (4NQO)-induced orthotopic OSCC tongue tumor, which shares 94% similarity with the human OSCC mutational landscape (14). To confirm its suitability, we first tested 4MOSC1 responses to Ser/Gly deprivation in vitro and observed a 60% reduction of cell proliferation in -Ser/Gly media (**Fig. S5A**), along with increased epithelial-like clustering and morphology (**Fig. S5B**), consistent with human patient-derived OSCC cells (**Fig. 1**; **Fig. S1**).

Following orthotopic injection of 4MOSC1 cells into the tongue, mice were maintained on a complete media diet for 6 days to allow tumor establishment, then randomized by sex to remain on either complete diet or switch to Ser/Gly-deprivation diet for 11 days. Body weights and tumor volumes were recorded every other day (**Fig. 7A**). PBS-injected mice on complete or Ser/Gly-deprivation diets served as negative controls. Tumor-free mice on Ser/Gly-deprivation diet showed no body weight changes or detectable tongue epithelial alterations, indicating no overt deleterious effects (**Fig. S5A** and **C, D**). Across tumor-bearing groups, body weight fluctuations were not significant from study initiation to completion (**Fig. S5B**), and Ser/Gly deprivation did not alter tongue epithelial histology (**Fig. S5E** and **F**). Tongue tumor measurements showed markedly reduced tumor growth within 24 hours of switching to the Ser/Gly-deprivation diet. Despite some fluctuation, tumors from Ser/Gly-deprived mice were 70% smaller than controls by day 16 post-injection, corresponding to 10 days after the dietary switch (**Fig. 7B** and **C**). Immunofluorescence imaging detected H3K27me3 and H3K4me3 expression throughout the tumors (**Fig. 7D** and **E**), although quantification did not show consistent changes in relative fluorescence for either mark.

**Figure 7.**
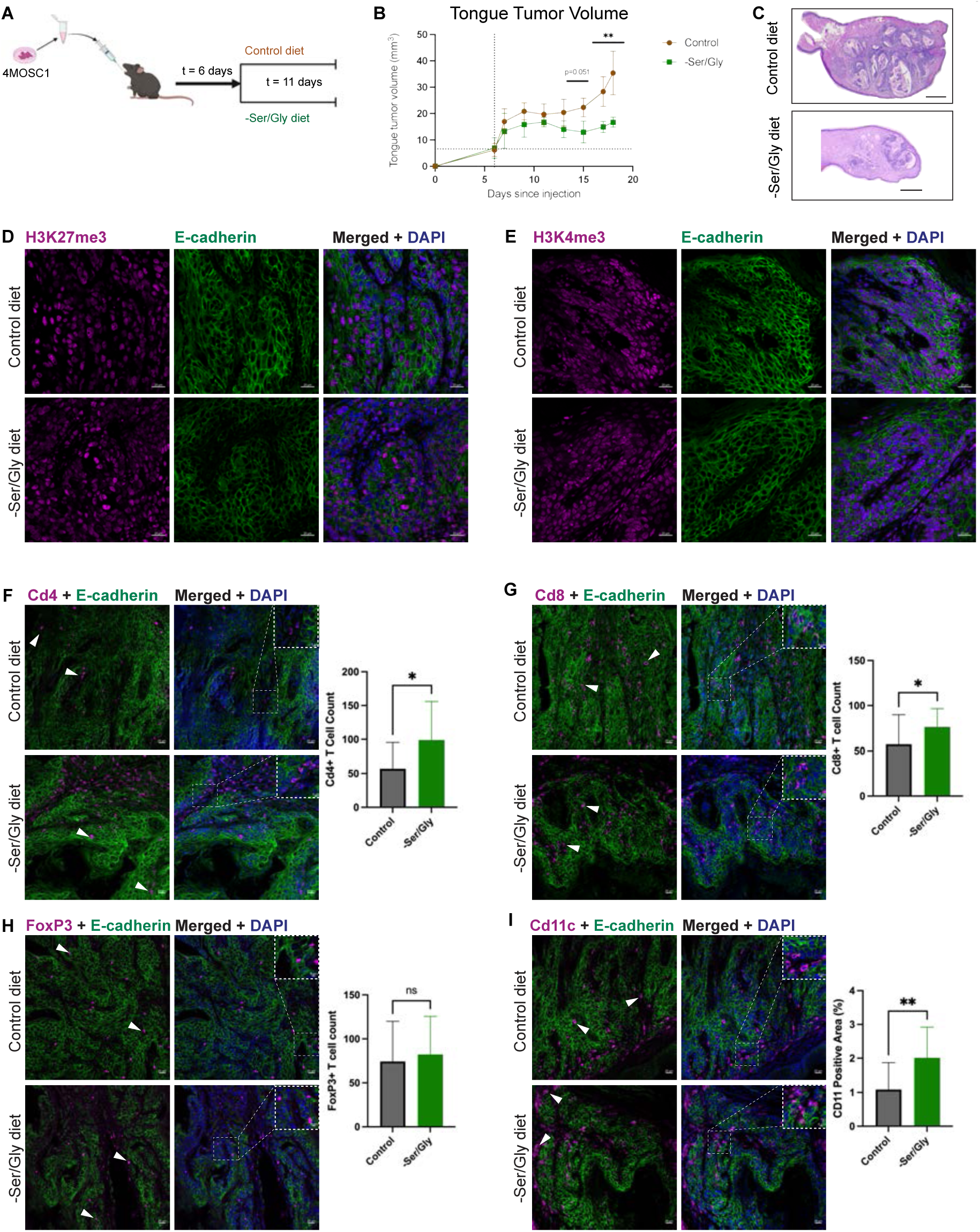
**A,** Schematic of in vivo mouse model experiment. 20-week-old C57BL6J male and female mice were injected with 1.0 x 10^6^ 4MOSC1 cells. Tumors were allowed to grow for 6 days before mice were either maintained on the control complete diet or switched to the Ser/Gly deprivation (-Ser/Gly) diet. Mice were maintained on respective diets for 12 days before sacrifice. Control complete media diet, n=4; Ser/Gly deprivation (-Ser/Gly) diet, n=5. **B,** Tongue tumor volumes were measured via caliper every other day. Tumors from mice on the Ser/Gly deprivation diet were 65% smaller than in mice on the complete, control diet. Vertical line at day 6 indicates time of diet change; all mice were given complete control diet prior to day 6. **C,** Representative H&E examples of mouse tongue tumors from the control complete diet and Ser/Gly deprivation diet; scale bar, 500 μm. **D,** Representative example and quantification of relative fluorescence unit (RFU) intensity of H3K27me3 of mouse tongue tumor from the control complete diet and -Ser/Gly diet. Scale bar, 20μm. Student’s unpaired t-test of average of 8 independent regions per mouse, n=3 control mice, n=4 -Ser/Gly mice. **E,** Representative example and quantification of relative fluorescence unit (RFU) intensity of H3K4me3 of mouse tongue tumor from the control diet and -Ser/Gly diet. Scale bar, 20μm. Student’s unpaired t-test of average of 8 independent regions per mouse, n=3 control mice, n=4 -Ser/Gly mice. **F** through **I**, Representative immunofluorescence of Cd4+, Cd8+, FoxP3+ and Cd11c+, immune cells in tongue tumors from mice on either the control, complete diet or mice on the Ser/Gly deprivation diet. Dotted boxes designate tumor areas and their corresponding magnified images. Arrowheads indicate immune cell staining in tumor areas. Scale bar, 100 μm.

Given the marked reduction in murine OSCC isograft growth after Ser/Gly deprivation, we performed an initial characterization of the tumor immune microenvironment 11 days after the dietary switch. Because Ser/Gly deprivation has been reported to alter immune landscapes in other cancer types (48), we examined immune infiltration by immunofluorescence. Tumors from Ser/Gly-deprived mice showed improved E-cadherin junctional organization and increased recruitment of CD4+ and CD8+ T cells, without an appreciable increase in Foxp3+ regulatory T cells (**Fig. 7F** and **H**, *arrowheads and magnified insets*). Ser/Gly deprivation also increased recruitment of CD11c+ antigen-presenting cells to tumors (**Fig. 7I**, *arrowheads and magnified insets*). Increased CD4+, CD8+, and CD11c+ immune infiltration suggests enhanced anti-tumor activity, likely contributing to reduced tumor growth. Together, these results demonstrate that dietary Ser/Gly deprivation suppresses OSCC growth in a syngeneic mouse model.

## Discussion

Studies presented in this manuscript provide insights into the complex relationship between nutrient metabolism, chromatin remodeling, and cell plasticity in OSCC. They present evidence that the removal of a non-essential amino acid, serine, has profound suppressive effects on OSCC cell proliferation and oncogenic traits in vitro and on the evolution of a murine orthotopic tongue tumor isograft. Additionally, they define a dual mechanism underlying the effects of Ser/Gly deprivation: the induction of SSP pathway with its rate-limiting enzyme, PSAT1, which produces αKG, the obligate substrate for KDM6B H3K27me3 demethylase, leading to induction of differentiation genes, and relocation of H3K27me3 to H3K4me3-marked promoters to generate bivalent chromatin domains that suppress aggressive cell states. Such a dual mechanism reverses cell plasticity with aggressive cell identities into more differentiated states suggesting their return to antecedent phenotypes. Collectively, our data indicate that under normal growth conditions OSCC cell identities are regulated by a reciprocal epigenetic program involving repression of differentiation genes and activation of stemness, EMT and cell cycle genes. Our work defines the major role of dietary Ser/Gly deprivation in suppressing aggressive OSCC cell states by activating a novel SSP-αKG-KDM-H3K4me3/H3K27me3 bivalency axis.

Under normal nutritional conditions with ample supply of dietary Ser/Gly, OSCC cells behave like serine auxotrophs, shutting down the endogenous SSP to maintain plastic cell states through the repressive chromatin mark H3K27me3 and activating H3K4me3. In response to Ser/Gly deprivation, OSCC cells upregulate the endogenous SSP enzymes, including PSAT1, leading to upregulation of αKG, activation of KDM6B/JMJD3 and demethylation of H3K27me3 (12). Indeed, we observed a decrease in the H3K27me3 repressive mark in two OSCC cell lines. Therefore, our data aligns increased αKG concentration, decreased H3K27me3 in immunoblots and immunofluorescence images along with a loss of H3K27me3 from differentiation genes’ loci in CUT & RUN-sequencing results and supports our conclusion that Ser/Gly deprivation in OSCC cells induces differentiation by via induction of KDM6B/JMJD3 activity.

Under Ser/Gly deprivation, the loss of H3K27me3 from its suppressive differentiation genes’ loci aligns with a concomitant reduction in H3K4me3 at promoters of stemness, proliferation and EMT genes. We find that this is associated with an establishment of genome-wide bivalency through relocation of H3K27me3 to the H3K4me3 promoters of genes driving aggressive cell states with no major change in H3K4me3 peak numbers. At this stage, it is unclear if the upregulation of SSP and production of αKG precedes the establishment of bivalency or if these mechanisms are implemented independently and converge to produce the overall outcome of de-repression of differentiation genes and inhibition of stemness, proliferation and EMT genes. Although our studies do not address the length of time of H3K27me3-H3K4me3-mediated bivalency, it suggests that the activation of the serine-αKG-KDM6B-bivalency axis via Ser/Gly deprivation may serve as a novel strategy to restrain OSCC evolution either alone or in combination with other therapies. We note that the observed prevalence of bivalency in OSCC cells highlights the importance of integrating epigenomic topography with gene expression, genetic variations and treatment-related environmental stresses to gain more profound understanding of their impact on epigenomic maps in OSCC and other cancer types.

Significantly, we find a reduction of plastic cell states under Ser/Gly deprivation conditions. There is a marked decrease in EMT and E2F target hallmark genes, both of which contribute to more aggressive tumors (4) in OSCC cell lines under Ser/Gly deprivation. The overexpression of *SERPINE1, TGM2*, and *LGALS1* is commonly associated with poor clinical outcomes and high risk of metastasis in HNC (42, 49, 50). Additionally, Ser/Gly deprivation is associated with upregulation of *mTORC1* repressors. Since our previous work has shown that mTORC1 signaling promotes p-EMT cell states in OSCC (45), targeting these plastic cell states via Ser/Gly deprivation may reduce tumors capacity to progress to advanced disease and/or improve response to therapy. Furthermore, our bulk RNA-seq data identified several aggressive hallmarks gene sets targeted by Ser/Gly deprivation treatment in OSCC cell lines. Therefore, the Ser/Gly deprivation signature may include useful biomarkers predictive of tumor sensitivity to the Ser/Gly deprivation diet.

Consistent with previous reports of a reduction in tumor growth under Ser/Gly deprivation diets in colon cancer (48), our in vitro results had in vivo significance. Mice given a Ser/Gly deprivation diet generated significantly smaller tumors with greatly reduced kinetics. We also found that tumors from mice on Ser/Gly deprivation diet displayed more focused E-cadherin junctional organization suggesting improved epithelial morphology consistent with downregulation of aggressive cell states. We note that while our in vitro data revealed reduced rates of proliferation, they reflected changes in cell states rather than cell death. At the same time, our vivo studies identifed increased presence of CD11c+ dendritic cells under Ser/Gly deprivation conditions, suggesting that the alteration of the chromatin landscape in OSCC may lead to the presentation of tumor antigens that trigger an immune response through antigen-presenting cells. Moreover, since epigenetic regulation plays central roles in immune cell lineages, their identities and activities, including T cells, the Ser/Gly deprivation diet is likely to impact their functions either independently of, or in concert with, epigenetic changes in OSCC epithelia.

Collectively, our studies provide mechanistic evidence supporting the anti-tumor effects of dietary Ser/Gly deprivation on serine metabolism in OSCC that, in turn, drive epigenetic changes in the chromatin landscape resulting in the induction of differentiation and reduction of aggressive cell states. Further investigation into how Ser/Gly deprivation diet drives changes in chromatin topology leading to the loss of cell plasticity in the tumor epithelia and how they impact the tumor stroma and the immune landscape is required to gain a detailed understanding of the relationships between the Ser/Gly deprivation diet and overall OSCC tumor biology. Such knowledge will be critical for translating dietary serine deprivation to the clinic to promote improved treatment strategies for OSCC patients.

## Acknowledgments

We acknowledge the BIDMC Mass Spectrometry Core Facility for their processing of our metabolomics samples and the DFCI Molecular Biology Core Facility for library preparation and sequencing of our CUT&RUN samples. We also thank the Boston University Microarray and Sequencing Resource Core Facility for processing our bulk RNA sequencing samples. The results shown here are in part based upon data generated by the TCGA Research Network.

## Data Availability

All CUT & RUN and RNA sequencing data have been deposited at GEO under accession number GSE286465 (Cut&Run) and GSE286464 (RNASeq). All code is available on github at https://github.com/montilab/SerineStarvation, which will be made available at publication.

## Authors’ Contributions

S.A.J. and M.A.K. conceptualized the project. S.A.J., L.K., M.V.B., X.V., S.M., and M.A.K. contributed to the methodology and experimental design. S.A.J., N.C.H., and M.V.B. performed experiments. S.A.J. and L.K. analyzed data and designed figures. S.A.J. and M.A.K. wrote the manuscript. All authors discussed results during the experimental process and contributed to editing the manuscript.

## Note

Supplementary data for this manuscript are included with manuscript resubmission

## List of Abbreviations

αKG: Alpha-ketoglutarate
CSC: Cancer stem-like cells
CUT&RUN: Cleavage under targets and release using nuclease
D2HG: 2 Hydroxyglutarate enantiomer D
EMT: Epithelial-to-mesenchymal transition
FRiP: Fraction of reads in peaks
GSEA: Gene set enrichment analysis
HNC: Head and neck cancers
JMJC: Jumonji domain C
KEGG: Kyoto Encyclopedia of Genes and Genomes
OSCC: Oral squamous cell carcinoma
PD-1: Programmed cell death 1
pEMT: partial-EMT
PHGDH: Phosphoglycerate dehydrogenase
PSAT1: Phosphoserine aminotransferase 1
PSPH: Phosphoserine phosphatase
-Ser/Gly: serine and glycine deprivation
SHMT1/2: Serine hydroxymethyltransferase ½
SMPDB: Small molecule pathway database
SSP: Serine synthesis pathway
TCGA: The Cancer Genome Atlas

