## Supplemental Figure Legends for "Dietary Serine Deprivation Impacts H3K27 and H3K4 Methyl Epigenomes to Impede Head and Neck Cancer Cell Plasticity and Tumor Growth"

**Supplementary Figures**

**Figure S1**. Ser/Gly deprivation inhibits proliferation of OSCC SCC25 and SCC9 cells and promotes epithelial phenotype. **A,** Proliferation of SCC25 cells grown in complete medium (black) was decreased by 58% when grown in Ser/Gly deprivation medium (-Ser/Gly, green) as determined by relative fluorescence units (RFU). Error bars represent ±SD. Student's unpaired t-test, n=3. **B,** SCC9 cells’ proliferation cultured in complete medium (black) was reduced by close to 90% when grown in Ser/Gly deprivation medium as determined by relative fluorescence units (RFU) (-Ser/Gly, green). Error bars represent ± SD. Student's unpaired t-test, n=3. **C,** Phase contrast imaging of SCC25 cells after 72 hours of growth in either complete or Ser/Gly deprivation media. Scale bar, 500 μm; inset scale bar, 100 μm. **D,** Phase contrast imaging of SCC-9 cells after 72 hours of growth in either complete or Ser/Gly deprivation media. Scale bar, 500 μm; inset scale bar, 100 μm.

**Figure S2.** Global metabolomics reveals limited changes in metabolites in Ser/Gly deprivation. **A,** Log_2_ -fold change in the standard 20 amino acids under -Ser/Gly conditions. Cells were grown in complete or Ser/Gly deprivation media for 48 hours. Analysis done using MetaboAnalyst. Blue bars indicate False Discovery Rate (FDR) p < 0.01. **B,** Volcano plot of all metabolites in -Ser/Gly media with FDR < 0.05 for both Log2 fold change ≤ 1(**blue**) and ≥ 1 (**red**). Analysis was done using MetaboAnalyst. **C,** Schematic of intermediate metabolites closely related to serine and glycine metabolism. Metabolites downregulated in -Ser/Gly media are shown in **blue**; metabolites upregulated in Ser/Gly deprivation media are in **red**. Metabolic pathways are highlighted in yellow. **D,** Enrichment of known Small Molecular Pathway Database (SMPDB) metabolite sets for altered metabolites in -Ser/Gly media.

**Figure S3.** Global RNA-seq reveals gene expression changes in OSCC CAL27 and HSC-3 cells grown in Ser/Gly deprivation media compared to growth in complete media. **A** and **B**, Heatmaps of indolent CAL27 and metastatic HSC-3 cells show statistically significant changes in the expression of genes under Ser/Gly deprivation conditions compared to complete media with more affected genes in metastatic in HSC-3 cells; n = 3 per cell line per condition. **C**, GSEA enrichment of HSC-3 cells-derived Ser/Gly RNA-seq gene signatures using squamous epithelial cell keratinization Human Protein Atlas geneset, with Normalized Enrichment Score (NES) of 1.5. **D** and **E**, CAL27-derived RNA-seq gene signatures were projected on The Cancer Genome Atlas (TCGA) OSCC RNA-seq database with known stage and grade and analyzed for association with OSCC progression. The Jonckheere–Terpstra (JT) test was used to determine monotonically increasing or decreasing relationships: for complete media, a positive JT value indicating that signature was increasing with stage/grade, whereas in Ser/Gly deprivation media, the negative value indicated that signature was decreasing with increased stage and grade.

**Figure S4. A,** CUT & RUN visualization of peaks overlapping LGALS1 gene. From top to bottom, samples are: Complete H3K4me3 (blue), -Ser/Gly H3K4me3 (green), Complete H3K27me3 (blue), -Ser/Gly H3K27me3 (green). Peak is most prominent in Complete H3K4me3, but also apparent in both -Ser/Gly histone marks. Peak is totally absent in Complete H3K27me3. **B,** CUT & RUN visualization of peaks overlapping SERPINE1 gene. From top to bottom, samples are: Complete H3K4me3 (blue), -Ser/Gly H3K4me3 (green), Complete H3K27me3 (blue), -Ser/Gly H3K27me3 (green). Peak is most prominent in Complete H3K4me3, but also apparent in both -Ser/Gly histone marks. Peak is totally absent in Complete H3K27me3. **C,** Full genome view of CUT & RUN peaks. Each row is a chromosome, with the x axis representing the chromosome coordinates. Within each chromosome row, there are four heatmaps corresponding to from top to bottom: -Ser/Gly H3K4me3, -Ser/Gly H3K27me3, Complete H3K4me3, and Complete H3K27me3. Color represents peak height at each location. -Ser/Gly H3K4me3, -Ser/Gly H3K27me3 and Complete H3K4me3 show overlapping peak regions, with Complete H3K4me3 showing overall greater peak intensity at most locations. The Complete H3K27me3 peaks are largely in different locations than the other three samples.

**Figure S5**. Serine deprivation diet has no detectable impact on mouse weight or tongue oral epithelia compared to control diet. **A,** Proliferation of 4MOSC1 cells grown in Ser/Gly deprivation medium (green) displayed decreased growth compared to cells grown in complete medium (black) as determined by relative fluorescence units (RFU). Error bars represent ±SD. Student's unpaired t-test, n=3. B, Phase contrast imaging of 4MOSC1 cells after 72 hours of growth in either complete or Ser/Gly deprivation media. Scale bar, 500 μm; inset scale bar, 100 μm. **C,** Male and female C57BL6J mice given a PBS injection showed no significant change in weight outside of normal fluctuations over 17 days. D**,** Male and female C57BL6J mice given 4MOSC1 injection showed no significant change in weight after change from complete diet at day 6 (vertical line) to Ser/Gly deprivation diet. E**,** H&E staining of healthy mouse with PBS injection given control diet for 18 days. Scale bar 500 µm; inset scale bar 100 µm. **F,** H&E staining of a mouse with PBS injection on Ser/Gly deprivation diet for 18 days. Scale bar 500 µm; inset scale bar 100 µm.
