## Supplementary figures and images for "Dietary Serine Deprivation Impacts H3K27 and H3K4 Methyl Epigenomes to Impede Head and Neck Cancer Cell Plasticity and Tumor Growth"

### Supplementary Figures

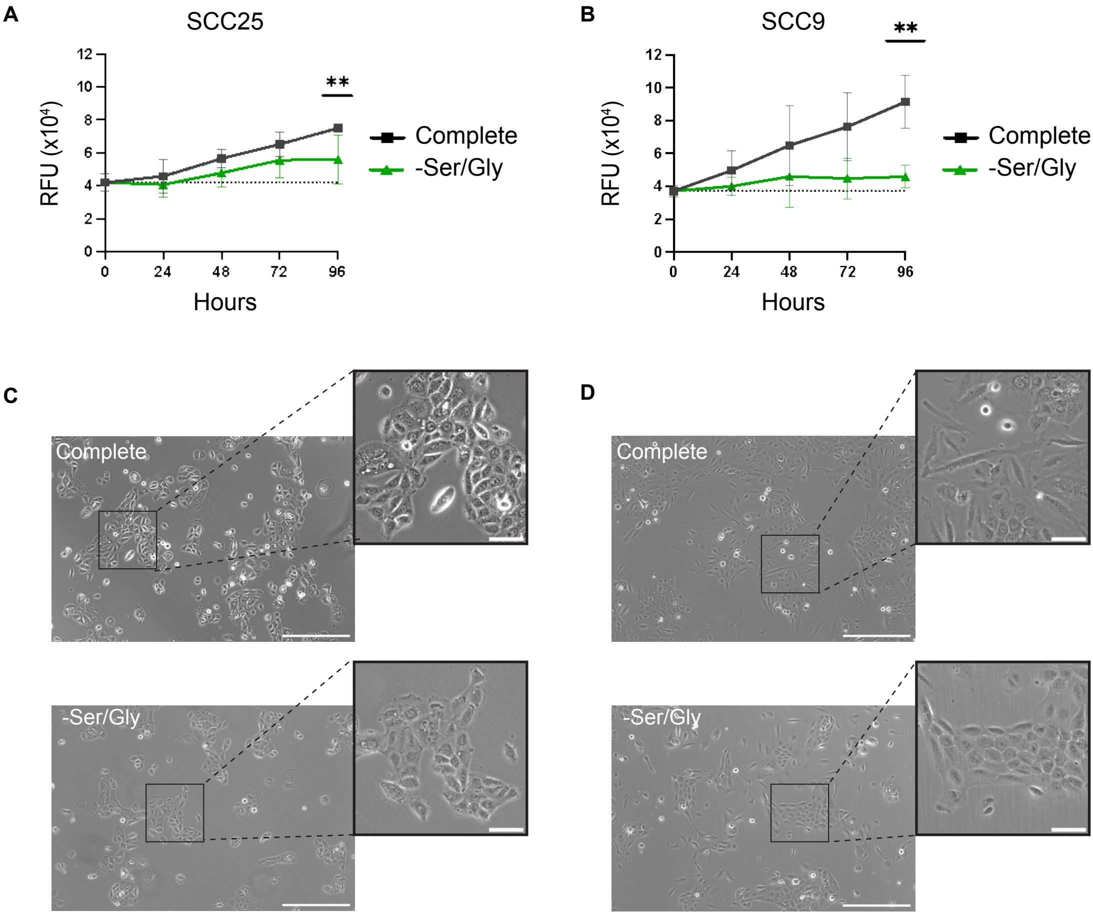

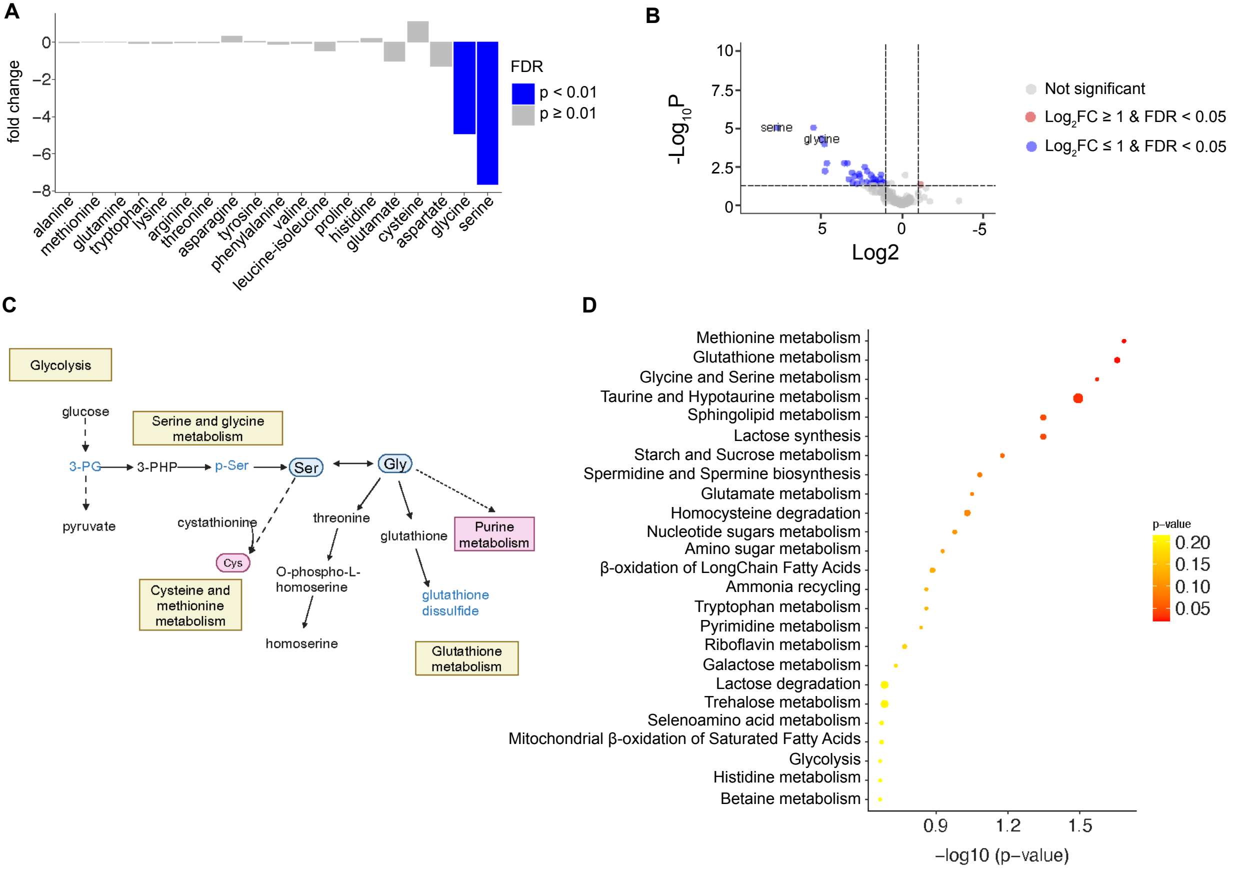

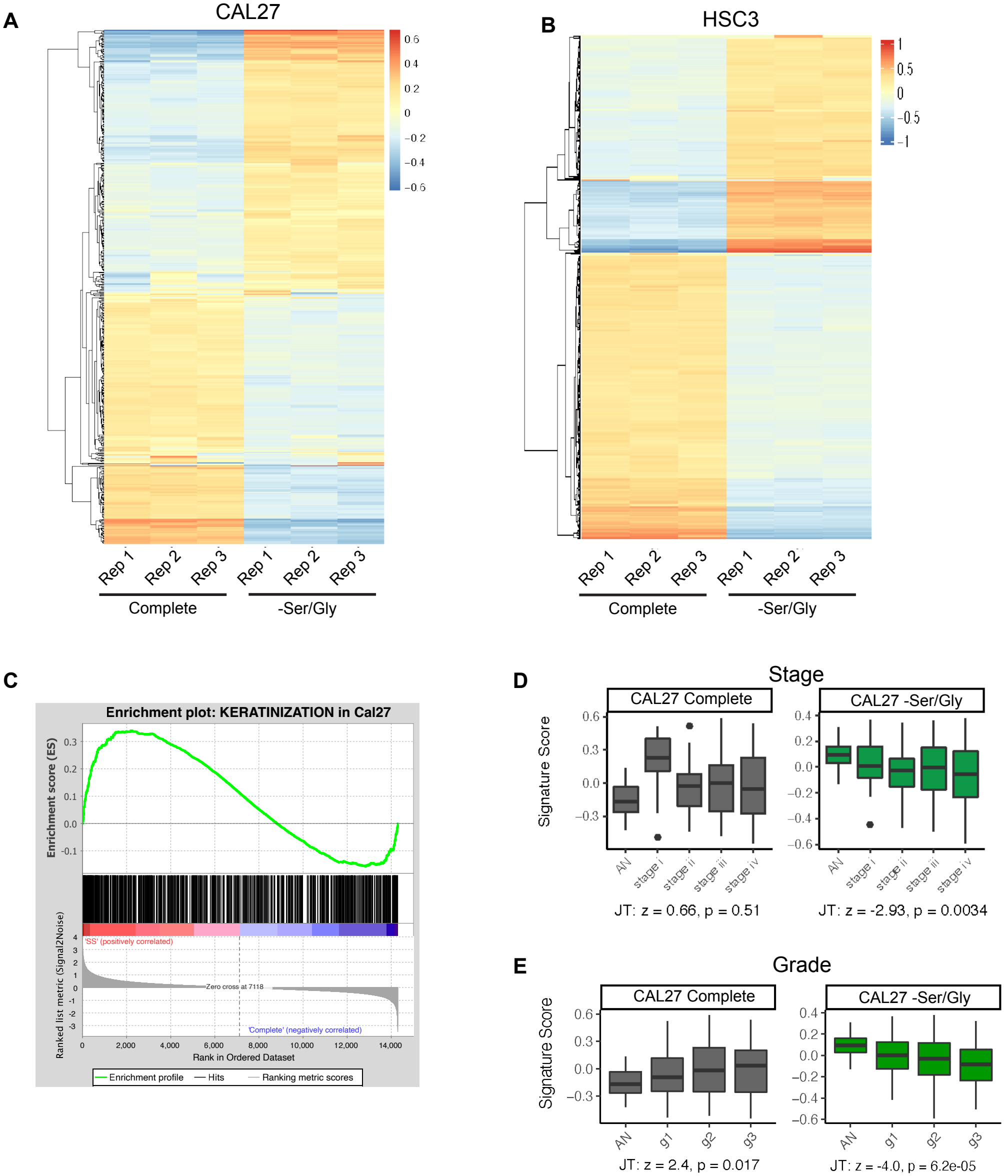

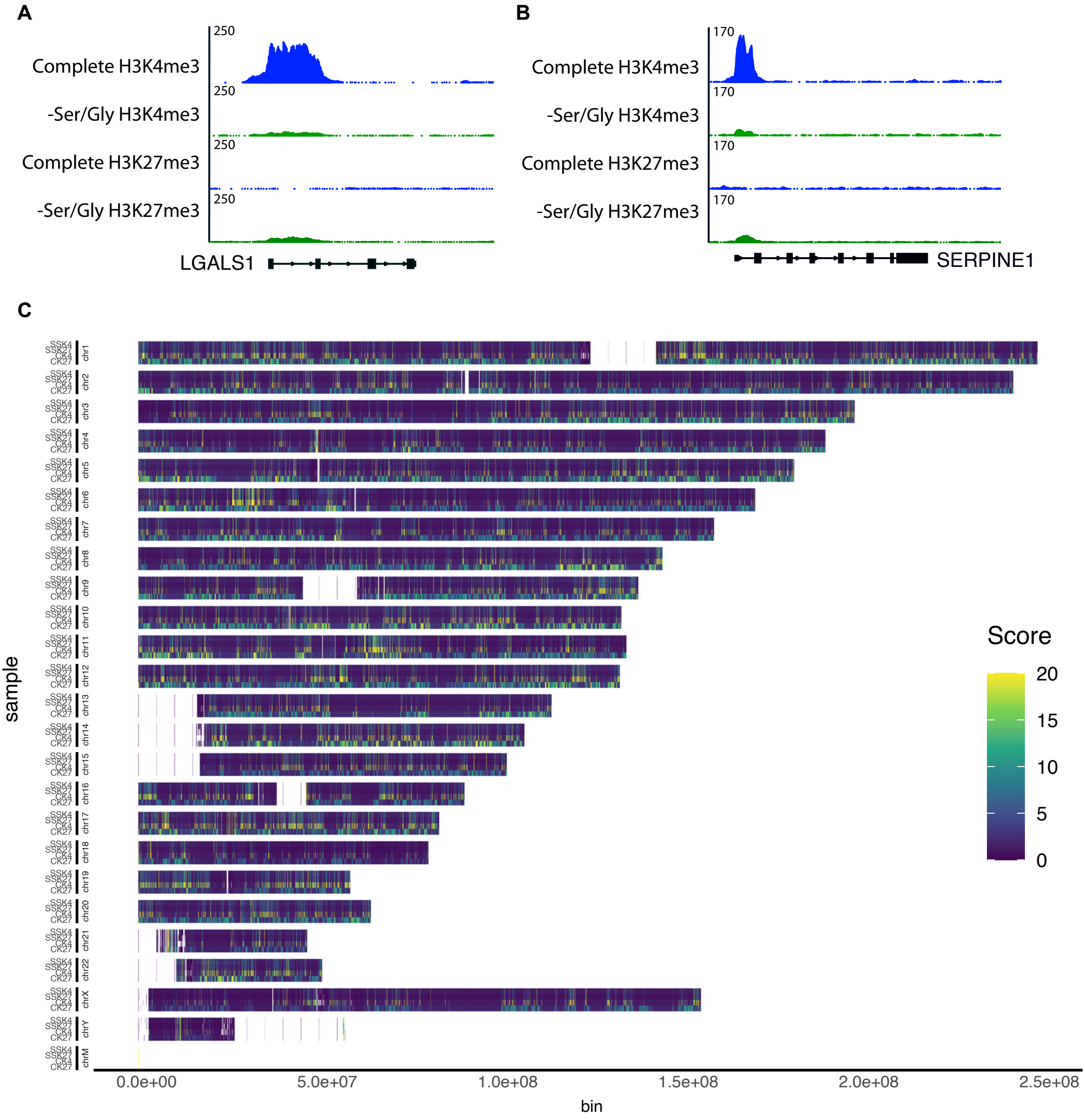

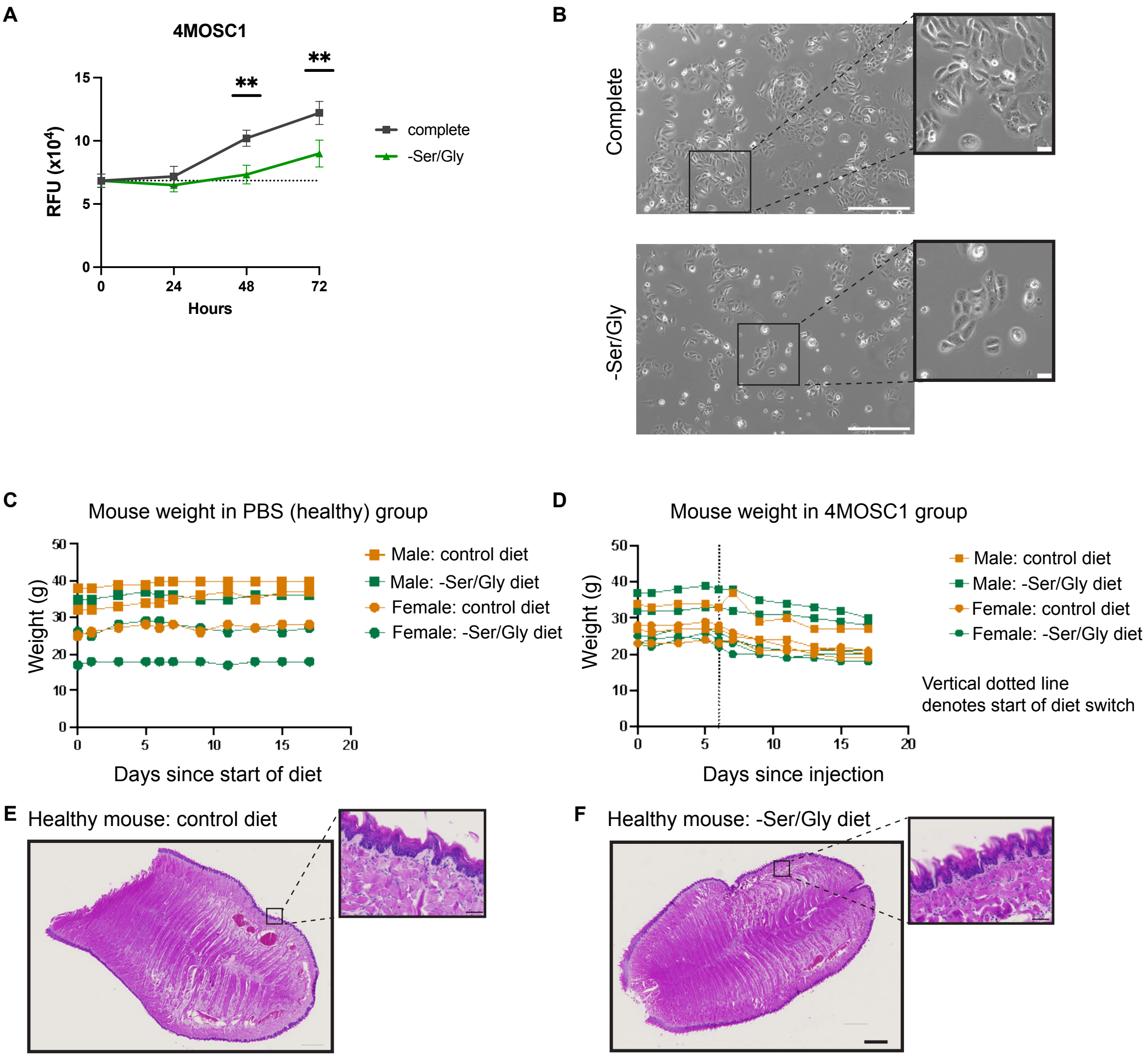
