## Supplementary Tables for "Dietary Serine Deprivation Impacts H3K27 and H3K4 Methyl Epigenomes to Impede Head and Neck Cancer Cell Plasticity and Tumor Growth"

|  | **CAL27** | | **HSC-3** | |
| --- | --- | --- | --- | --- |
| **Gene name** | **log2FC** | **FDR** | **log2FC** | **FDR** |
| PHGDH | 1.234 | 6.03E-164 | 0.840 | 1.46E-112 |
| PSAT1 | 0.994 | 2.24E-145 | 1.503 | 4.03E-296 |
| PSPH | 0.607 | 3.00E-34 | 0.502 | 1.44E-23 |

**Table S1.** Gene Expression of serine synthesis pathway enzymes is significantly increased in both CAL27 and HSC-3.

| **RT-qPCR primer** | **Sequence** | **Source** |
| --- | --- | --- |
| PHGDH fwd | ATCTCTCACGGGGGTTGTG | Maddocks et al., 2013^28^ |
| PHGDH rev | AGGCTCGCATCAGTGTCC | Maddocks et al., 2013^28^ |
| PSAT1 fwd | CGGTCCTGGAATACAAGGTG | Maddocks et al., 2013^28^ |
| PSAT rev | AACCAAGCCCATGACGTAGA | Maddocks et al., 2013^28^ |
| PSPH fwd | CGGTCCTGGAATACAAGGTG | Maddocks et al., 2013^28^ |
| PSPH rev | CAGGGAGGTGAGCTGTGC | Maddocks et al., 2013^28^ |
| Actin fwd | TCCATCATGAAGTGTGACG | Maddocks et al., 2013^28^ |
| Actin rev | TACTCCTGCTTGCTGATCCAC | Maddocks et al., 2013^28^ |

**Table S2.** List of primers used in this study. Related to STAR Methods.

|  | **CAL27** | | **HSC-3** | |
| --- | --- | --- | --- | --- |
| **Gene name** | **log2FC** | **FDR** | **log2FC** | **FDR** |
| SERPINE1 | -0.0743 | 0.655 | -1.720 | <1E-300 |
| COL12A1 | -0.6379 | 9.74E-83 | -2.222 | <1E-300 |
| TGM2 | -0.7790 | 1.80E-16 | -1.404 | <1E-300 |
| IGFBP3 | -0.7187 | 7.36E-84 | -2.161 | 4.63E-201 |
| LGALS1 | -2.0805 | 2.79E-154 | -1.573 | 1.91E-181 |
| SPARC | -1.0378 | 8.38E-82 | -1.110 | 2.24E-138 |
| VIM | -1.6052 | 4.99E-03 | -0.787 | 3.61E-133 |
| MATN2 | -0.8679 | 2.99E-40 | -1.547 | 3.01E-113 |
| CDH2 | -0.9219 | 1.20E-37 | -1.433 | 1.88E-70 |

**Table S3.** Select EMT genes show decreased expression in CAL27 and HSC-3 cells in serine deprivation conditions.

| **Sample** | **Total Reads** | **RiP** | **FRiP** |
| --- | --- | --- | --- |
| **IgG Control** |  |  |  |
| IgG_1 | 59045967 | 1234954 | 0.0209 |
| IgG_2 | 49954181 | 2792307 | 0.0558 |
| IgG_3 | 44180220 | 828021 | 0.0187 |
| **H3K27me3** |  |  |  |
| **Complete_H3K27me3_1** | 93212848 | 54473720 | 0.5844 |
| **Complete_H3K27me3_2** | 49195179 | 6947179 | 0.1412 |
| **Complete_H3K27me3_3** | 57490846 | 4632237 | 0.0805 |
| **-Ser/Gly_H3K27me3_1** | 78929908 | 6572930 | 0.0832 |
| **-Ser/Gly_H3K27me3_2** | 46737636 | 13134494 | 0.281 |
| **-Ser/Gly_H3K27me3_3** | 47685220 | 6303362 | 0.1321 |
| **H3K4me3** |  |  |  |
| **Complete_H3K4me3_1** | 57491939 | 37343006 | 0.6495 |
| **Complete_H3K4me3_2** | 50200189 | 595122 | 0.0118 |
| **Complete_H3K4me3_3** | 36126091 | 100944 | 0.0027 |
| **-Ser/Gly_H3K4me3_1** | 68210104 | 23414519 | 0.3433 |
| **-Ser/Gly_H3K4me3_2** | 49209111 | 10254107 | 0.2083 |
| **-Ser/Gly_H3K4me3_3** | 36126091 | 3966206 | 0.1098 |

**Table S4.** Fraction of reads in peak (FRiP) scores of CUT&RUN replicates to determine how many reads fall into peaks within each sample. Samples with a FRiP score greater than 0.1 were included in further analysis.
